# Progressive community, biogeochemical and evolutionary remodeling of the soil microbiome underpins long-term desert ecosystem restoration

**DOI:** 10.1101/2023.09.26.559499

**Authors:** Qiong Chen, Mengyi Yuan, Liuyiqi Jiang, Xin Wei, Zhen Liu, Chen Peng, Zinuo Huang, Dongmei Tang, Xiangrong Wu, Jing Sun, Cunqi Ye, Qing Liu, Xiaowei Zhu, Peng Gao, Laibin Huang, Meng Wang, Mingkai Jiang, Chao Jiang

**Author notes:** These authors contributed equally (Qiong Chen, Mengyi Yuan, Liuyiqi Jiang, Xin Wei). These authors contributed equally (Zhen Liu, Chen Peng, Zinuo Huang, Dongmei Tang). Senior author (Qiong Chen).

## Abstract

Ecological restoration of degraded lands is essential to human sustainability. Yet, an in-depth community, functional, and evolutionary microbial perspective of long-term restoration of damaged ecosystems is lacking. Herein, we comprehensively assessed the impact of long-term (up to 17 years) restoration of Tengger Desert, China, by multi-omic profiling of 1,910 topsoil samples. The soil biophysiochemical properties, especially soil hydraulics, microbiome stability, and functional diversity, significantly improved during restoration. The soil microbiome transitioned from an extreme oligotrophic and autotrophic community to a diverse copiotrophic ecosystem. The soil microbiota, including fungi, could mediate the soil physicochemical changes through metabolites. Importantly, the systematic rewiring of nutrient cycles featured the multi-domain preference of an efficient carbon fixation strategy in the extreme desert environment. Finally, the microbiome was evolving via positive selections of genes of biogeochemical cycles, resistance, and motility. In summary, we present a comprehensive community, functional, biogeochemical, and evolutionary landscape of the soil microbiome during the long-term restoration of desert environments. We highlight the crucial microbial role in restoration from soil hydraulic and biogeochemical perspectives, offering promising field applications.

**Highlights:**

- The desert soil microbiome transformed from simple oligotrophic to a diverse, stable, and nutrient-rich ecosystem with expanded functional diversity.
- Restoration led to systematically rewired biogeochemical cycles, which are highly efficient in carbon fixation in the desert environment.
- The microbiome was evolving via positive selections of genes involved in biogeochemical cycles and environmental adaptations.
- Microbes and metabolites could facilitate desert restoration from hydraulic and biogeochemical aspects, offering promising field applications.

## Introduction

Ecological restoration is pivotal in combating land degradation and ensuring a sustainable future for all^1^. Land degradation of drylands, comprising about 41% of the world’s land area and habitats to 38% of the global population^2,3^, is defined as desertification. Global restoration efforts, such as the Great Green Wall projects in Africa and China^4^, and society-sponsored restoration efforts, such as Alxa Tengger Desert Lockside Ecological Project and the Alipay Ant Forest Project, all contribute to desert ecosystem restoration^5^. Encouragingly, satellite data (2000-2017) revealed a striking increase in vegetation in China and India, with China accounting for 25% of the global net increase in leaf area^6^.

Vegetation restoration is effective for rehabilitating degraded lands and slowing desertification in arid regions^7–9^. The microbiome is essential in soil and plant functioning^10^. Studies have shown that soil- and plant-associated microbiomes could function as biofertilizers and biocontrol agents to enhance soil hydraulic properties^11^, fertility, plant stress tolerance, and productivity^12–15^. Studies on desert metabolites focused on plant- and endophytes-derived metabolites, which are important for environmental adaptation and drug discovery^16,17^. However, fewer studies focused on the dynamics and functional role of soil microbiome during long-term desert ecosystem restoration. Previous research has investigated the changes in the carbon, nitrogen^18,19^, hydraulic properties^20^, and variations in the community composition and function of nematodes^21^ and fungi^22^ during restoration. We hypothesized that the microbiome directly contributes to and responds to desert ecological restoration from pan-domain, functional, ecological, biogeochemical, and evolutionary aspects. A detailed understanding of the role of microbial components is essential for developing an effective ecological restoration approach.

In this study, we collected 1910 topsoil samples from experimental sites of up to 17 years of restoration at the eastern edge of the Tengger Desert (Methods), located south of the Gobi Desert. Different planting (polyculture vs. monoculture) and irrigation strategies (drip and manual subsurface irrigation) were implemented at different sites. We modified our protocols for studying air microbiome^23,24^ to overcome the low-abundance issue in desert microbiome^25^. We integrated microbiome and metabolome data with soil physiochemical properties to show that soil health and ecosystem stability gradually improved at the polyculture site with traditional sub-surface irrigation. The microbiome could mediate soil health through metabolites and nutrient enrichments. Importantly, the microbiome showed the systematic rewiring of biogeochemical cycles and an increasing repertoire of secondary functions over time. Finally, the biogeochemical genes, among others, were evolving to adapt to the changing environment. We provide an integrated physicochemical, compositional, ecological, functional, biogeochemical, and evolutionary perspective on the role of soil microbiome during long-term desert restoration, with promising field applications.

## Results

### Desert soil biophysiochemical properties improved during restoration

In brief, 1910 samples were collected from two restoration sites within a week in 2020 (Table S1). Of which, 1490 samples (pooled into 298 testing samples) were from a polyculture restoration site (Figures 1A, 1B, S1A-S1C, and Table S1; Methods); 420 samples (pooled into 84 testing samples) were collected from a monoculture restoration site (Figures S1A, S1D, S1E, and Table S1). The polyculture site implemented two irrigation methods: manual subsurface irrigation (SI) and drip irrigation (DI). SI requires drilling a hole for each plant and watering 3 to 5 times/year for the first three years, while DI involves dripping water from a piping system that waters indefinitely. Nearby natural sand dunes (SD) were collected as the negative control (Figure 1A). The sub-groups of SI and DI regions were named based on the duration of restoration (Figure S1C). The DI regions had more visible vegetation (Figure S1C).

**Figure 1.**
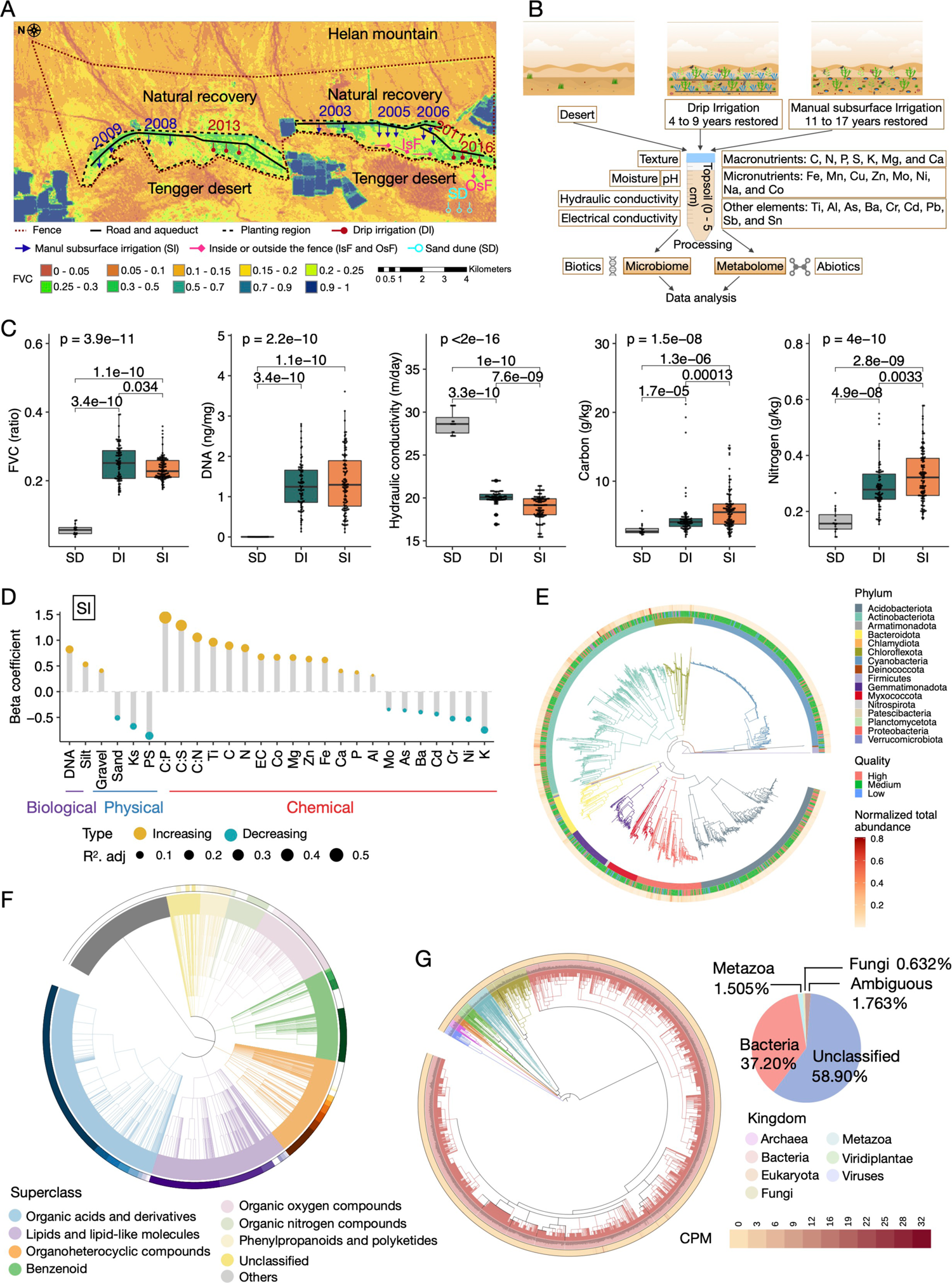
Integrated multi-omic profiling of soil microbiome during long-term desert restoration. (A) Fractional vegetation cover (FVC) index of the sampling sites at the polyculture site. The numbers of samples in each group were SD (17), OsF (22), IsF (53), DI (83), and SI (123). (B) The multi-omic experimental design. (C) Representative differentially abundant soil biophysiochemical properties. (D) Effects of the restoration duration on soil health properties at the SI regions (GLM; *p.adj* < 0.05). The beta coefficient represents the directions and magnitudes of the restoration effects. PS, particle size; Ks, hydraulic conductivity; FVC, fractional vegetation cover. (E) Phylogenetic trees of bacterial metagenome-assembled genomes (MAGs). The branches and inner ring are colored by Kingdom. The intermediate ring denotes high (red), medium (green), and low (blue) quality MAGs. The outer ring indicates the relative abundance of MAGs. (F) Total diversity of metabolome. The colors of the outer ring denote different superclasses. (G) Total diversity of the microbiome. The pie chart indicates the total relative abundance of respective kingdoms. The outer ring of the tree indicates the relative abundance of CPM. See also Figures S1 and S2.

At the polyculture site, restoration with SI improved the soil biophysiochemical properties, as demonstrated by significantly increased (Kruskal-Wallis and Generalized Linear Model [GLM]; *p.adj* < 0.05) soil DNA abundances, fractional vegetation cover (FVC), hydraulic properties (1/Ks), electrical conductivity (EC), macronutrients (C, N, P, Mg, and Ca), micronutrients (Fe, Zn, and Co), and other helpful elements (Ti and Al; Figures 1C, 1D, S1F, and S2A)^26,27^. Still, a significant gap exists between the recovered nutrient level and the arable topsoil (Figure S2A)^28,29^. The DI method was not as effective but showed significantly increased (*p.adj* < 0.05) DNA abundance, FVC, and hydraulic properties over time (Figures 1C and S2B). These results demonstrated that soil health attributes, including hydraulic properties, nutrient levels, and biological abundances, improved gradually during restoration.

At the monoculture site, the soil biophysiochemical properties didn’t significantly improve (Figure S2C). Non-biocrust (NBC) and biocrust (BC) samples were collected (Figures S1D and S1E). The natural recovery (NR) area was generally covered by biocrusts.

Nearby sand dunes (SD2) were used as the negative control (Figures S1D and S1E). As expected, the NBC group was more similar to the SD2 control, and the BC group was more comparable to the NR group (Figure S2D). BC and NR groups had higher hydraulic properties, carbon, and DNA concentrations (Figure S2D). Overall, we show that biocrusts are important for water retention and can improve carbon accumulation in soil, consistent with previous findings^30^.

In conclusion, restoring the desert ecosystem is a gradual but measurable process. Soil biophysiochemical properties significantly improved over time at the polyculture site.

### The diverse desert soil metagenome-assembled genomes and metabolome

Each sample was sequenced by the Illumina NovaSeq 6000 platform, resulting in a total of 28.5 billion reads (8.74 TB) for 382 samples, with an average of 149 million paired-end reads (22.88 GB) per sample (Methods; Figures S2E-S2G). To assess contamination, we incorporated blank controls in the experimental procedures, from which no data could be generated (Methods). We assembled 431.0 million contigs (Table S1), with 585.7 million genes predicted, and binned them into 1565 nonredundant Metagenome-Assembled Genomes (MAGs; Methods), of which 300 MAGs were high-quality (≥90% completeness and <5% contamination) (Figures S2H-S2J).

The 1565 MAGs were classified by the Genome Taxonomy Database Toolkit (GTDB-Tk) ^31^ (Figures 1E and S2I) and organized into species-level genome bins (SGBs) at an ANI threshold of 95%, resulting in a total of 710 prokaryotic SGBs. Strikingly, most (96.8%) of SGBs were not similar to existing species and were defined as unknown SGBs (uSGBs). The 687 uSGBs were phylogenetically diverse, with Actinobacteriota (Actinobacteria) and Acidobacteriota (Acidobacteria) dominating (Figure S2J). Notably, the proportion of uSGBs in most phyla was close to 100%, drastically different from the results of recovering MAGs from human metagenomes^32^, indicating our limited knowledge of desert microbiomes.

For microbial metabolic activity, we profiled the metabolites with liquid chromatography (C18 column) coupled with tandem mass spectrometry in both positive and negative electrospray ionization modes. We obtained 26,055 features, among which 21,486 had MS^2^ information. After data cleaning and feature extraction (Methods), we obtained 18,885 features. 1,449 features were matched with comprehensive metabolome databases (including NIST, GNPS, and MoNA; Table S2; Methods), and 17,436 were annotated using SIRIUS/CSI: FIngerID (Figure 1F). Still, the low annotation rate (1449/18885) underscores a pressing need to expand the knowledge of desert microbial metabolome.

In short, we generated a large collection of new desert soil MAGs and metabolome datasets, and there is still much to be discovered in the biological and chemical diversity in the desert soil ecosystems.

### Desert microbiome dynamics indicate that the soil ecological systems transformed from simple and oligotrophic to complex and copious

To gain a more comprehensive understanding of the composition of the soil microbiota, as the GTDB database did not cover non-prokaryotes, we established an extensive reference genome database covering more than 431,109 species across all domains of life (Methods). We used Kraken2^33^ to classify 41.4% of the total reads (Figure 1G) and hyperbolic arcsine transformed counts per million (aCPM) values for downstream analyses unless noted^34^ (Methods). In total, we identified 59 phyla, 1287 genera, and 4344 species (Figure 1G).

We first focused on the dynamics of SI regions of the polyculture site as they showed more systematic improvements with longer restoration duration. The Shannon indices of bacteria and fungi in the SI regions increased over time (Figure S3A). Notably, the SD region had the lowest Shannon indices of bacteria and fungi but the highest Shannon indices of archaea and viruses (Figure S3A). Principal Coordinate Analysis (PCoA) based on species profiles showed a significant separation of SD, SD2, and NBC groups from the others (Figure 2A). Notably, the two control sites, SD and SD2, clustered together (Figure 2A). The NR group was clustered with the BC group, away from the polyculture regions (Figure 2A). The IsF group was more clustered with the SI regions than the OsF group, as IsF samples were collected inside the fence of the SI regions (Figure 1A). The microbiome composition transformed along the duration gradient in the SI regions (Figures S3B and S3C). At the phylum level, the top seven phyla plus unidentified bacteria and metazoan accounted for 98.06% of the relative abundance across all samples (Figure 2B) at the polyculture site. Actinobacteria (46.4%), Proteobacteria (19.0%), and Cyanobacteria (5.09%) dominated (Figure 2B), consistent with previous studies^12,13^.

**Figure 2.**
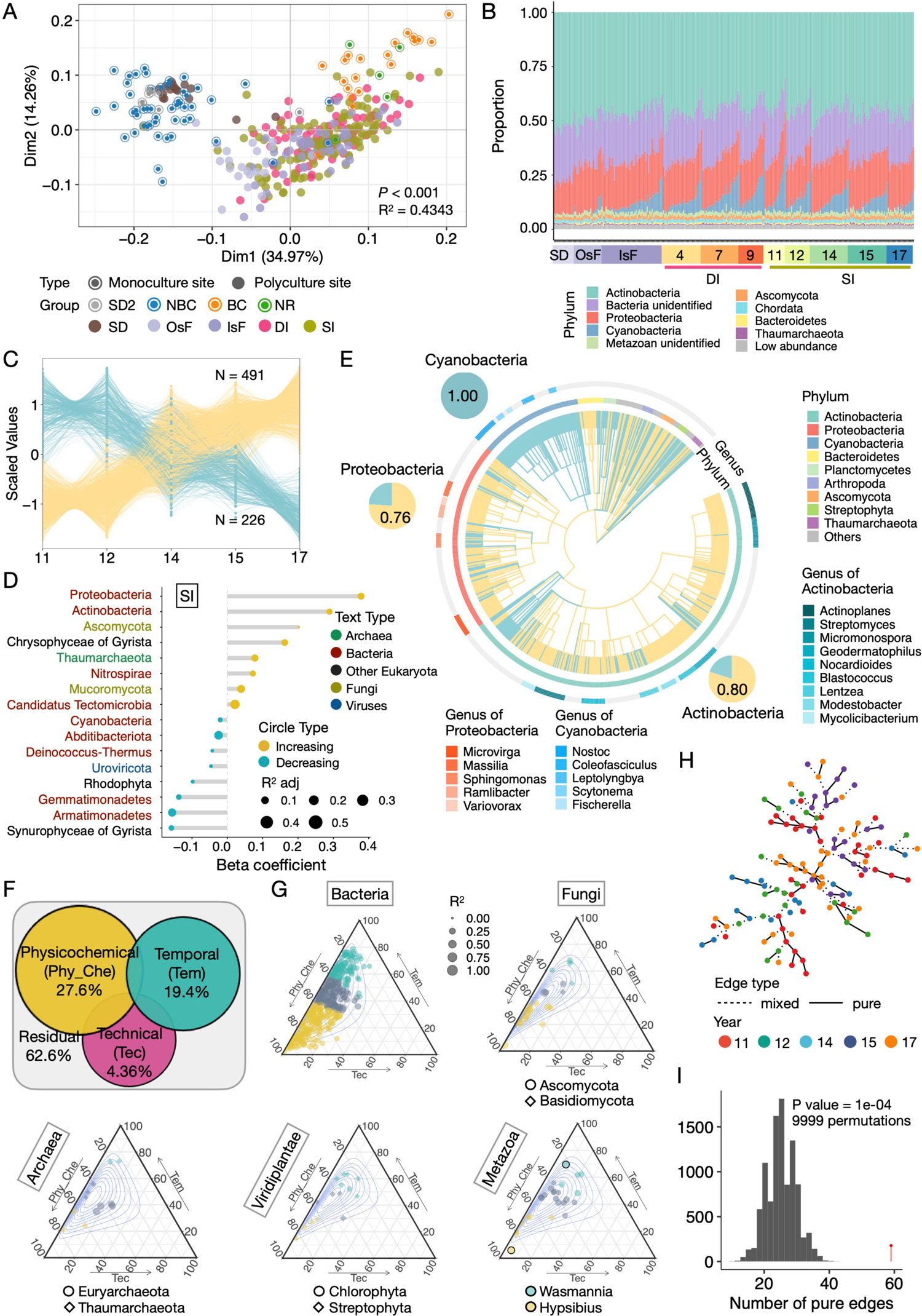
The sources of variation of the highly dynamic desert soil microbiome. (A) PCoA analysis of all samples at the species level. Dots represent samples. Colors and styles represent sampling groups and sites. (B) The relative abundance of the top 10 phyla at the polyculture site. (C) The increasing (yellow) and decreasing (green) clusters of the species abundance profiles in the SI regions, membership > 0.8. (D) Regression-based analysis of temporal variation of phyla in the SI regions (GLM; *p.adj* < 0.05). (E) The dynamically changing microbes in the SI regions. Only species with temporal patterns estimated by GLM (*p.adj* < 0.05), spearman correlation (*p.adj* < 0.05), or C-means clustering (membership > 0.8) were shown. The colors of the branches in the inner ring correspond to increasing (yellow) or decreasing (blue) species. The second and third rings correspond to phyla and genera. Pie charts reflect the relative proportions of increasing and decreasing species in a phylum. (F) Partial dbRDA variation-partitioning analysis of SI samples. (G) Ternary plots of variation-partitioning analysis of SI samples. Each dot represents a genus; the size corresponds to the total explained variation. Physiochemical (dark yellow) and temporal (blue) variables can play dominant roles in species distribution, or neither (gray). Contours denote 0.1 to 0.9 confidence intervals. (H) The Nearest Neighbor (NN) tree of the SI regions. Nodes are connected by solid edges (pure) if from the same group. Otherwise, dashed lines were used (mixed). Colors denote the duration of restoration. (I) Graph-based permutation test (N=9999) on the NN tree generated from (H), red pin denotes the observed number of pure edges. See also Figures S3 and S9.

The 1430 species identified from the SI regions were separated into two clusters by fuzzy c-means (Figure 2C). GLM analyses (*p.adj* < 0.05) showed that Proteobacteria, Actinobacteria, Ascomycota, Chrysophyceae (golden algae), Thaumarchaeota, Nitrospirae, Mucoromycota, and Candidatus Tectomicrobia increased over the years (Figure 2D). For example, the top increasing Proteobacteria genera were members of rhizosphere plant-microbes, such as symbiotic nitrogen-fixing *Microvirga*^35^, antibiotics and plant hormone producer *Massilia*^36^, and desiccation-resistant *Ramlibacter*^37^(Figures 2D and 2E). The top increasing Actinobacteria genera included producers of antibiotics: *Actinoplanes*^38^, *Micromonospora*^39^, and *Streptomyces*^39^, followed by a gamma-radiation-resistant genus *Geodermatophilus*^40^ (Figures 2D and 2E). The two fungal phyla Ascomycota and Mucoromycota are key drivers of the arable soil decomposition process^41,42^; The Nitrospirae and archaeal Thaumarchaeota participate in nitrogen cycles^43,44^.

In contrast, Armatimonadetes, Deinococcus-Thermus, Abditibacteriota, Cyanobacteria, and other phyla decreased over time (Figure 2D). These microorganisms are highly resistant and can survive in oligotrophic habitats. For example, the photoautotrophic prokaryotes Cyanobacteria inhabit oligotrophic arid environments^45^. Armatimonadetes strains are aerobic oligotrophs^46^; Abditibacteriota is oligotrophic and extremely resistant to antibiotics and toxic compounds^47^. The dynamically changing pan-domain phyla support the gradual shift from an oligotrophic to a more copiotrophic environment in the SI regions over time.

### The sources of variation in the desert soil microbiome dynamics

The extensive microbiome dynamics prompted us to identify the sources of variation in the desert soil microbiome during restoration. Compositional dissimilarities (beta diversity) were partitioned into species replacement and richness components (Methods). Species replacement processes dominated bacterial (73.6%), fungal (52.9%), and metazoan (62.3%) communities (Figure S3D). The contribution of richness difference was higher for archaea (73.7%).

Physicochemical (Phy_Che), temporal (Tem), and technical (Tec) variables, such as technical batch effects, can impact the soil microbiome composition. To evaluate the temporal influence rigorously, we constructed asymmetric eigenvector maps (AEM) variables to extract broad- or fine-scale temporal profiles (Figure S3E; Methods). 46 meta-variables (including 8 AEM variables) were collected and classified into physiochemical, temporal, and technical groups. We performed partial distance-based redundancy analysis (dbRDA) to decompose the variation in the SI regions after variable selection (Figures S3F and S3G). The temporal, physiochemical, and technical variables explained 19.4%, 27.6%, and 4.36% of the variation (squared Bray-Curtis distance) in the SI soil microbiome (Figures 2F and S3G). Together, the selected variables explained 37.4% of the soil microbiome variation (Figure S3G).

We next employed multivariate regression-based variation partitioning analysis on the 565 genera in the SI regions to estimate the effects on individual genera. After filtering genera that were invariable (*p.adj* ≥ 0.05), the 449 genera were used to perform hierarchical partition analysis to evaluate the relative importance of each group (Methods). These genera covered bacteria (371/449), metazoan (26/449), fungi (22/449), and viridiplantae (16/449). We define a group of variables that explained more variation than the others (1.50-fold) as dominating. For bacteria, 96 genera were subjected to dominating temporal influences, while 149 were subjected to dominating physicochemical influences (Figure 2G). Cyanobacteria were subjected to more dominating temporal (7/18) than physiochemical influences (1/18) (Figure 2G). In addition, 5 archaea, 22 fungi, and 11 viridiplantae genera were subjected to dominating physiochemical or temporal influences (Figure 2G). Chlorophyta, or green algae, was dominated by temporal variables (Figure 2G). No genera were subjected to dominating technical influence, indicating the robustness of our methods.

To further investigate whether samples of the same duration of restoration were more similar, we constructed the nearest neighbor (NN) tree (Figure 2H; Methods)^48^. The graph-based permutation test (N = 9999) showed that pure edges were enriched (*p* = 0.0001) in the SI regions, indicating that samples from the same group were more similar (Figures 2H and 2I). We also constructed a time-predictive model with the soil microbial profiles from the SI regions (Figures S3H-S3K; Methods). The model was stable with a median multi-class area under the curve (mAUC) of 0.718 (Figure S3I).

In short, the desert soil microbiome dynamics were influenced by physiochemical and temporal variables during long-term restoration, and different genera across domains were subjected to diverging influences.

### SI enhances the microbial community stability at the polyculture site

We assessed the microbiome assembly dynamics using methods developed based on neutral-based and niche-based theories^49^. The two methods showed that stochastic processes drove bacterial community assembly in the SD and SI regions (Figures 3A, S4A, and S4B; Methods). Then, we constructed species interaction networks at the polyculture site to systematically evaluate the ecological network dynamics during long-term restoration (Methods). The pan-domain network (Figure 3B) consisted of 1744 nodes (1591 bacteria, 53 metazoans, 37 fungi, 33 archaea, and others) and 40630 edges (92.57% positive and 7.43% negative). The SI regions showed less interdependence in the community (Figure S4C). The SD region showed drastically different network characteristics in almost every category (Figure S4C). Keystone taxa, playing crucial roles in a community (Methods), were identified from the SD (3) and SI (28) regions, which belonged to Proteobacteria, Euryarchaeota, and Thaumarchaeota (Figure S4D). These results demonstrated that archaea are critical in community structure besides bacteria. Importantly, the SI network became more stable over time, as estimated by increased robustness (*p* = 0.051) and cohesion (*p* = 4.09e-09) (Figure 3C). We employed a multiple regression model to reveal the detailed effects of biotic and abiotic properties on network stability (Methods). Abiotic and biotic variables together explained 90.5% of the variance in network stability (Figure 3D). All biotic variables positively impacted stability. Some abiotic variables, such as Ti and C:P ratio, positively impacted stability, while C: N ratio, Mn, and Ks negatively impacted stability.

**Figure 3.**
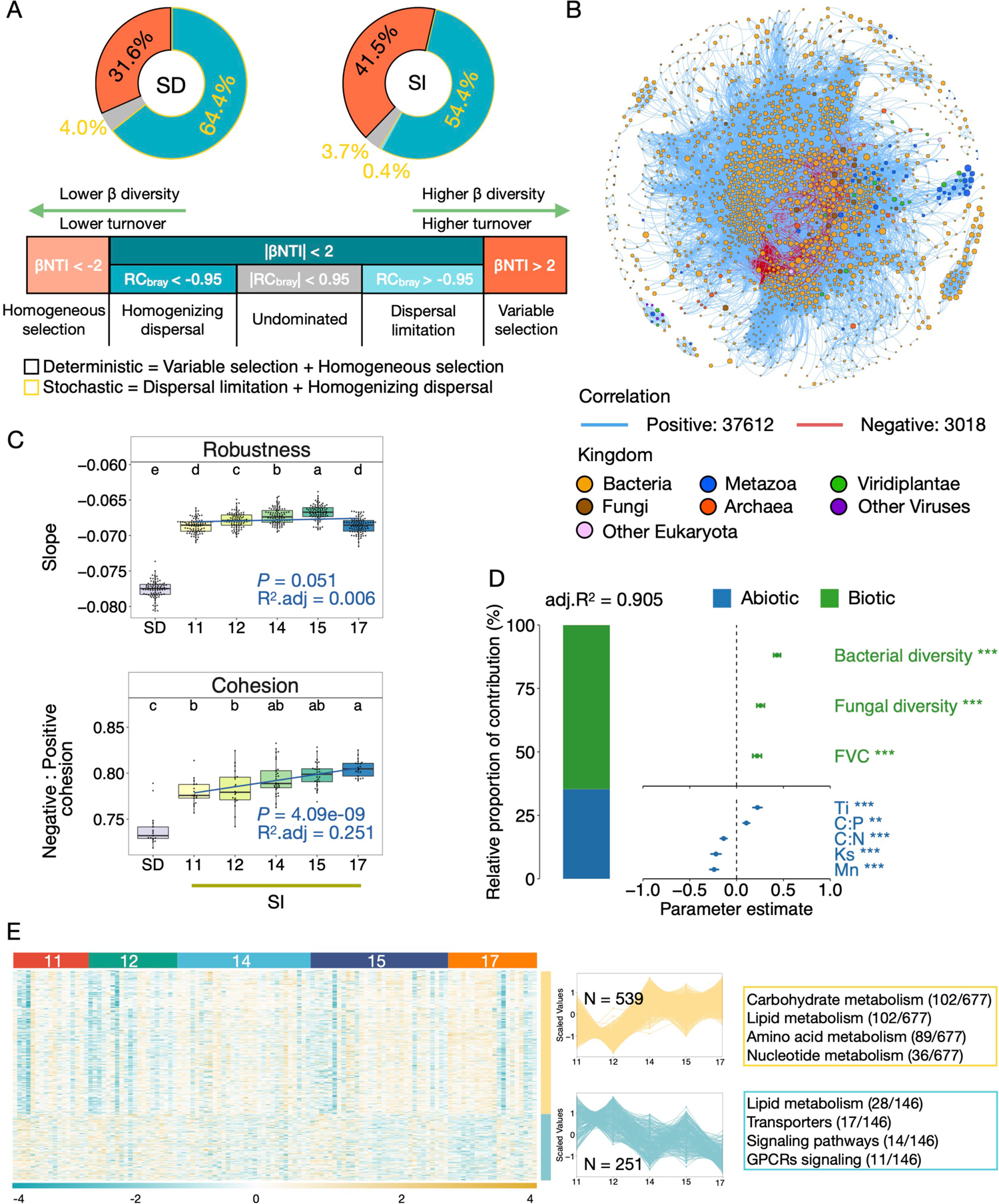
The dynamic microbiome community assembly processes, stability, and metabolites. (A) Pie charts map the proportions of ecological processes. βNTI: between-community nearest taxon index; RC_bray_: Bray–Curtis-based Raup–Crick. (B) The interaction network of pan-domain species at the polyculture site. Node colors represent different kingdoms. Node sizes indicate relative abundance (aCPM). Edges are colored based on positive (teal) and negative (red) interactions. (C) Community stability at the polyculture site. Different lowercase letters above boxplots indicate *p* < 0.05 (Kruskal-Wallis). The GLM statistics are also shown. (D) The relative proportions of the contribution of soil biotic and abiotic factors on cohesion in the SD and SI regions. The averaged standardized regression coefficients for the predictors are displayed with 95% confidence intervals. \*\*\**P* < 0.001; \*\**P* < 0.01; \**P* < 0.05. (E) Heatmap of increasing (yellow) and decreasing (green) metabolites based on C-means clustering (membership > 0.75) and GAM (*p* < 0.03) in the SI regions. The top four functional categories were labeled. See also Figures S4 and S10.

### Improved and diversified desert metabolome during restoration

We analyzed the metabolic functional diversity at the polyculture site by constructing a global network of metabolomic features (with MS^2^) based on the potential bio- or synthetic-relationships. The network had 5483 nodes and 16,969 edges representing biotic and abiotic relationships (Figure S4E). Among these, 75.36% of features were unclassified, showing an urgent need for further research (Figure S4E). Biotransformation accounted for 21 relationships, including methylation, deamination, and hydroxylation (Figure S4F), revealing diverse biochemical reactions that link the metabolite network. Organoheterocyclic compounds, organic acids and derivatives, and lipids and lipid-like molecules dominated (Figure S4G). 1449 annotated metabolites from SI samples were clustered into two groups, with 539 increased and 251 decreased over time (Figures 3E and S4H). The top increased metabolites analyzed by GAM (*p* < 0.03) provide energy for microorganisms and plants, exhibit antimicrobial properties, and play vital roles in plant-microbe interactions^50^ (Figure S4I). The top decreased metabolite phthalic anhydrides disrupt the balance of the soil microbial community and soil fertility^51^(Figure S4I). Metabolites with antimicrobial activities also decreased, such as kaurene diterpenoids^52^(Figure S4I). The increasing cluster was mainly related to carbohydrate, lipid, amino acids, and nucleotide metabolism (Figure 3E), essential for organismal growth. The decreasing cluster was mainly related to lipid metabolism, transporters, signaling pathways, and GPCRs signaling (Figure 3E). Notably, the number of lipid metabolism-related features in the increasing cluster were higher than that of the decreasing cluster (Figure S4J).

We further constructed AEM variables to extract temporal profiles. We found that AEM1 had the lowest *p*-value and increased over time, suggesting that AEM1 represents the major temporal trend of metabolome during restoration (Figure S4K). Overall, the dynamic metabolome patterns suggest that the soil is becoming more nutritious and life-supporting during restoration.

### Desert microbial functional transformation during restoration

To determine how the functional capacity of the soil microbiota developed during restoration, we annotated the genes from SI samples using the KEGG database (KOs; Methods). In total, 9734 KOs were mapped to 263 KEGG pathways. Of these, 133 pathways increased, and 97 pathways decreased over time (membership > 0.65, Figure 4A), also supported by regression analyses (145 pathways *p.adj* < 0.05; Figure 4B). We multiplied the relative abundance by the DNA concentration as a proxy of absolute abundance and found that all pathways gradually increased over time and peaked at 15 years (Figure S5A). Next, we used relative abundance to evaluate the functional diversity of the microbiome. In the SI regions (Figures 4B and 4C), the relative abundance of catabolic carbohydrate, lipid metabolism, and amino acid metabolism pathways increased, consistent with the metabolomic analysis (Figure 3E). Conversely, the abundance of anabolic carbohydrate and fatty acid pathways decreased over time. Interestingly, xenobiotic biodegradation and metabolism pathways, such as bisphenol, caprolactam, and dioxin degradation, increased while antibiotic biosynthesis decreased over time. Biofilm formation^53^ also decreased over time, consistent with the decreasing phyla adapted to oligotrophic environments (Figure 2D). Quorum sensing and bacterial chemotaxis increased, important in motility, symbiosis, and stability^54^. We further compared the pathways enriched in regions with the shortest (11 years) and longest (17 years) restoration duration in the SI regions by Reporter Score (Methods), with similar results (Figure S5B). Overall, the relative anabolic and antibiotic functions decreased, while the overall catabolic functions and functional diversity increased over time in the SI regions.

**Figure 4.**
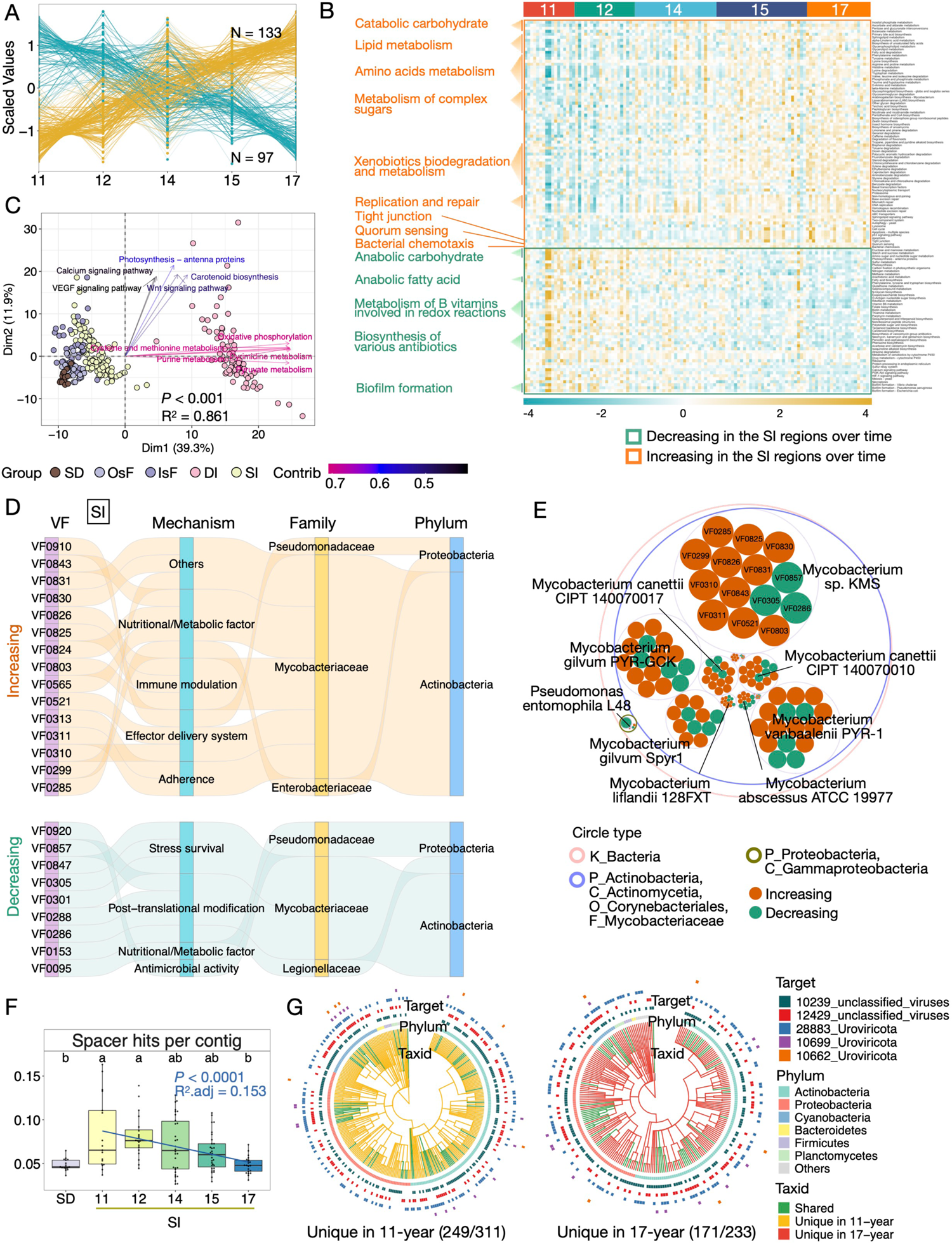
The transforming functional diversity of desert soil microbiota. (A) Fuzzy c-means clustering of the increasing (yellow) and decreasing (green) KEGG pathways in the SI regions. (B) Heatmap of significantly differed KEGG pathways of the SI regions (GLM; *p.adj* < 0.05). Representative functions are labeled on the left. (C) PCA bi-plot of all KEGG pathways at the polyculture site. Colored arrows denote the relative importance of contributing features. (D) Sankey diagrams showing the linkages between VFs, VF categories, and corresponding taxa at the family and phylum level for the increasing (top) and decreasing (bottom) VF profiles in the SI regions (GLM; *p.adj* < 0.05). (E) Bacteria producing diverse VFs (GLM; *p.adj* < 0.05) in the SI regions. The open circles are colored by taxonomic information. The filled circles represent temporally increasing (red) and decreasing (green) VFs. Circle (grey) size indicates the species-level average CPM relative abundance; representative species were labeled. (F) Spacers density changes over time in the SD and SI regions. The GLM statistics were labeled. (G) Hosts and targets of spacers in the 11-year and 17-year groups. Phylogenetic trees represent the host species encoding spacers. Branches are colored by whether the taxa are unique (yellow or red) in each group or shared (green). The inner ring indicates the phylum, and the outter rings indicate the taxonomy of the spacer targets. See also Figures S5 and S10.

The abundant antibiotics-related pathways promoted us to annotate 2,143 antibiotic resistance genes (ARGs) in the polyculture site, conferring resistance to 40 types of antibiotics. The most abundant classes were antibiotic inactivation (65%) and target alteration (17%; Figure S5C). The most abundant ARGs were carbapenem resistance genes (23%), followed by monobactam (18%) and cephalosporin (14%) resistance genes (Figure S5D). However, different regions did not show significant differences in PCA analysis (Figure S5E).

Virulence factors (VFs) undermine competitors. 848 VFs were annotated in the polyculture site. VFs belonged to 13 categories, with nutritional and metabolic factors (20.3%), effector delivery system (18.1%), and adherence (16.9%) dominating (Figure S5F). PCA analysis separated the SD control and other groups (Figure S5G). The SD region showed more VFs with adherence and post-translational functions than the SI regions (Figure S5H). Different organisms were involved in VF production for the SD control and the 17-year group. The diversity of VFs and producing organisms also increased during restoration (Figures S5I-S5K).

Ten VFs categories showed significant temporal patterns in the SI regions. Stress survival and antimicrobial VFs decreased; adherence, exotoxins, and nutritional/metabolic VFs increased over time (Figure S5I). Of the 767 VFs annotated in the SI regions, 26 increased, and 14 decreased over time (Figures 4D and S5M). The physicochemical properties, including pH, Ti, N, Mn, and C: N ratio, were related to the VFs patterns in the SI regions (Figure S5N). The family Mycobacteriaceae and the class Gammaproteobacteria produced the most dynamic VFs (Figures 4D and 4E). Notably, *Mycobacterium* sp. KMS encoded 14 VFs, involved in diverse functions such as post-translational modification, stress survival, and metabolism (Figures 4D and 4E). These results suggest that VFs are an important species-species interaction modulating mechanism during restoration.

Bacteria and archaea use CRISPR systems to defend against foreign DNA invasion^55^. At the polyculture site, 54,251 CRISPR arrays and 204,909 spacers were identified, and 60% of spacers were unique (98% similarity; methods). The spacers per contig decreased in the SI regions over time (Figure 4F). Actinobacteria, Proteobacteria, and Cyanobacteria dominated the CRISPR-hosts against five groups of viruses (hits range from 20 to 989) in 11-year and 17-year groups (Figure 4G). However, CRISPR hosts were mostly unique in each group, indicating dramatic species successions over time (Figures 4G and 3A). These results suggest that the CRISPR-Cas defense system became less utilized during restoration, implicating that the pressure of foreign DNA/RNA invasion had subsided over time.

To summarize, functional pathways, metabolites, VFs, and CRISPRs were involved in species and environmental interactions and significantly changed over time, supporting a more diversified and interactive microbial lifestyle during restoration.

### Remodeling of carbon metabolism during restoration

Carbon fixation is a critical step in restoring degraded lands. We derived six autotrophic CO_2_ fixation pathways from KEGG annotation results (Figures 5A, S6A, and S6B). In the SI regions, the Hydroxypropionate-hydroxybutyrate (HP/HB; e.g., *abfD*, *ACAT*, and *MUT*) cycle increased, while the Calvin cycle (e.g., *rbcL* and *rbcS*) and Wood-Ljungdahl pathways (e.g., *metF* and *folD*) decreased over time (Figure 5A). The demand for metals, cofactors, and anaerobiosis restricts the Wood-Ljungdahl pathway to extreme ecological niches^56^. The HP/HB cycle requires one-third less energy than the Calvin cycle in ammonia-oxidizing bacteria^57^ and does not have as high demands for metals and coenzymes as the Wood-Ljungdahl pathway^56^, and has potentially rapid metabolic pathway kinetics^58^. Additionally, its preference for bicarbonate (rather than CO_2_) makes it applicable in slightly alkaline environments. The HP/HB cycle also has excellent thermotolerance and oxygen tolerance^56^, making it well-suited for desert environmental conditions. Proteobacteria, Actinobacteria, Bacteroidetes, Firmicutes, and Cyanobacteria were the top phyla responsible for changes in carbon fixation (Figures 5A-5D and S6C). Interestingly, archaeal and fungal genes participating in carbons cycles also consistently changed to HP/HB cycle over time (Figures 5B-5D, bottom). These variations demonstrate that microorganisms are important founding autotrophs in the desert ecosystem, and the carbon fixation strategies significantly shifted during restoration.

**Figure 5.**
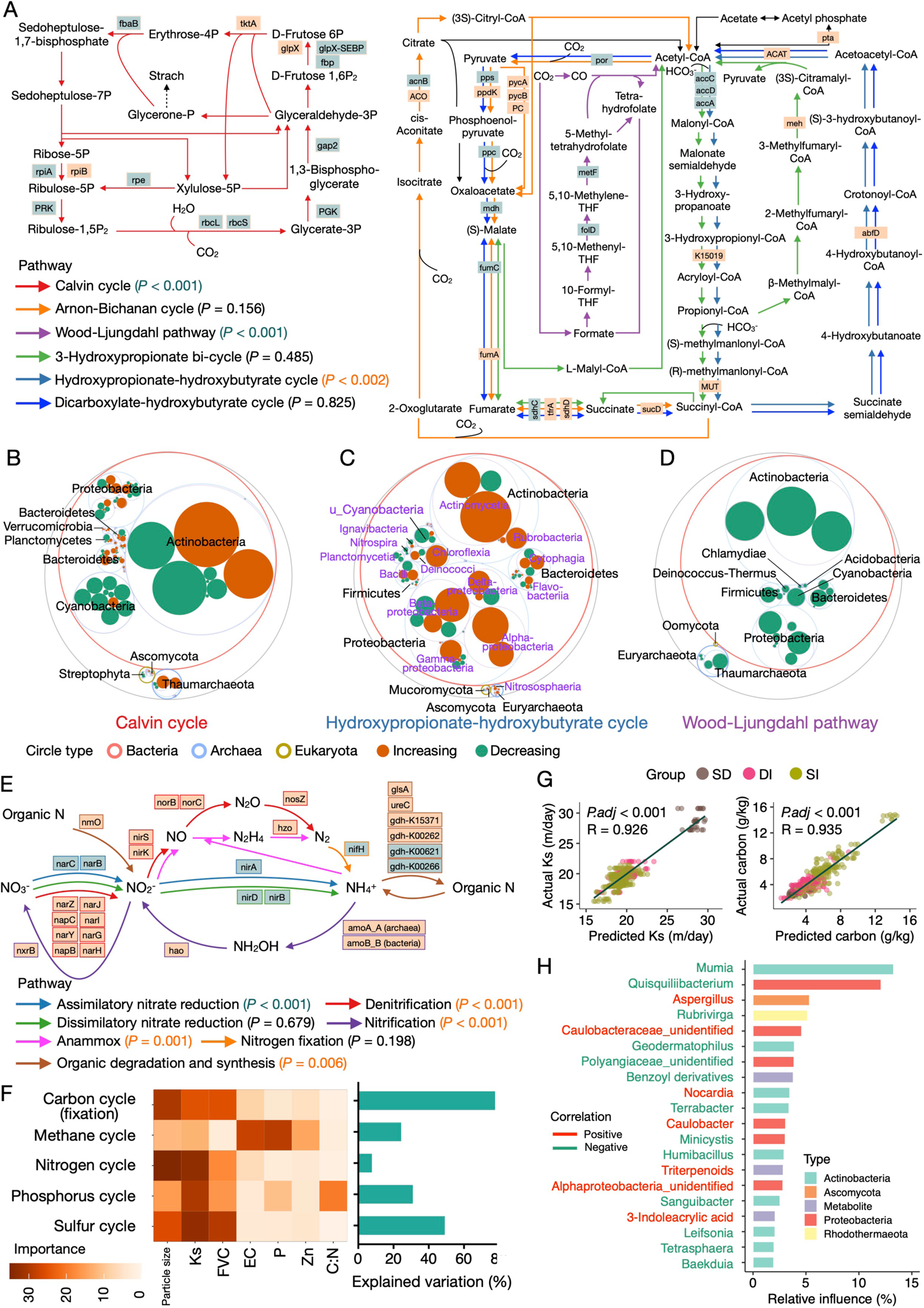
The systematically rewired biogeochemical cycles and predictive modeling of Ks and carbon in the SI region. (A) Six autotrophic CO_2_ fixation cycles. The shaded colors of enzymes denote temporally increasing (yellow) and decreasing (blue) enzymes (GLM, *p.adj* < 0.05). *P*-values estimated by GLM are displayed for each pathway with increasing (yellow) and decreasing (blue) patterns (B-D) The integrated taxon-function maptree of microbiome members involved in the Calvin cycle (B), Hydroxypropionate-hydroxybutyrate cycle (C), and Wood-Ljungdahl pathway (D). Outmost to innermost circles indicate the cellular organisms, kingdom, phylum, and class levels. The filled circles represent enzymes, as labeled in (A), that are temporally increasing (red) or decreasing (green). Circle size indicates the taxon-level average relative CPM abundance in the SI region. The phyla (black labels; light blue circles) with the top relative abundance are labeled. The classes (purple labels; light purple circles) with the top relative abundance are labeled in (C). (E) The simplified nitrogen cycle diagram, color-coded as in (A). (F) Contributions of soil health properties to nutrient cycles in the SD and SI regions. Colors in the heatmap represent variable importance calculated via multivariate linear regression modeling. The bar plots indicate the total explained proportions of the variation for each nutrient cycle. (G) Correlations between predicted and actual Ks (left) and carbon (right) as modeled by the gradient boosting regression models. (H) The top 20 variables for predicting the carbon level. Text color indicates positive (red) or negative (green) correlations with the carbon level as determined by GLM. See also Figures S6, S7, and S8.

Besides fixation, the other primary carbon source is degradation. We identified 349 carbohydrate-active enzymes (CAZYmes) in the SI regions^59^. Glycoside hydrolases (GHs, 46.4%) and glycosyltransferases (GTs, 34.0%) were the most abundant CAZYme families (Figure S6D). In the SI regions, the relative abundance of catabolic glycoside hydrolases (GHs), carbohydrate esterases (CEs), and polysaccharide lyases (PLs) increased; anabolic glycosyltransferases (GTs, formation of glycosidic bonds) decreased over time (Figure S6D), again supporting an increasing relative catabolic and a decreasing assimilative ability of soil microbial organisms during restoration (Figures 4B and S5B). We further characterized the enzymes responsible for degrading specific carbon substrates, including cellulose, hemicellulose, xylan, and lichenin. The relevant enzymes increased in the SI group (Figures S6E and S6F). Notably, an oversized increase in enzymes degrading lichenin was observed in archaea (Figure S6E). Even viruses may degrade cellulose and xylan, although the relative proportions were much smaller (Figure S6E). These results suggest that carbon degradation was increasingly utilized during restoration.

Methane (CH_4_) is one of the carbon sources and greenhouse gas. We reconstructed the CH_4_ cycle based on the MCycDB^60^ and annotated 225 genes. Methanogenesis converts methanogenic substrates to CH_4_ and is classified into three types — hydrogenotrophic, aceticlastic, and methylotrophic methanogenesis. Hydrogenotrophic methanogenesis uses H_2_ and reduces CO_2_ to CH_4_ *via* the Wood-Ljungdahl pathway^61,62^; the reduced H_2_ levels lead to acetate formation^62^, which is cleaved by aceticlastic methanogenesis to form CH_4_ and CO_2_. Methylotrophic methanogenesis dominates in sulfate-rich and saline systems^62^. In the SI regions, hydrogenotrophic (*hmd* and *ftr*) and methylotrophic (*torC* and *mttC*) methanogenesis increased over time, with methylotrophic methanogenesis dominating (Figures S7A-S7F). For consumption of CH_4_, both anaerobic (e.g., *narH*, *narG*, and *narZ*) and aerobic (28 involved genes increased) oxidation pathways increased in the SI regions (Figure S7A). These results indicate that CH_4_ is an important carbon and energy source for soil microbes and that microorganisms play a key role in maintaining the balance of CH_4_ in the desert environment. Further analyses showed that the genes of related pathways also increased, and major methanogenesis organisms included Planctomycetes, Proteobacteria, Actinobacteria, and Acidobacteria (Figures S7D and S7E). Notably, the presence of methylotrophic methanogenesis of Ascomycota origin was observed (Figure S7E).

In the SD region, the lowest abundance of anaerobic oxidation of CH_4_ (Figures S6A and S6B) coincided with the decreased denitrification and sulfur reduction (Figure S6B), likely decreasing available SO_4_^2-^and NO_x_^-^ as electron acceptors^63–65^. Hydrogenotrophic methanogenesis was activated to create conditions for substantial methanogenesis (Figure S6B), consistent with the enhanced Wood-Ljungdahl pathway (Figure S6B).

In summary, CO_2_ and CH_4_ are important carbon sources for desert soil microbiome, and microorganisms from bacterial, archaeal, fungi, and even viral domains all contribute to systematically rewiring carbon metabolic pathways during restoration.

### Systematic rewiring of biogeochemical cycles in the restoring desert soil ecosystems

In addition to carbon, nitrogen, sulfur (Supplemental information; Figures S7G-S7M), and phosphorous cycles (Supplemental information; Figures S8A-S8E) are essential in global biogeochemistry. We annotated 66 genes in the nitrogen cycle with the NCycDB^66^. In the SI regions, the relative abundance of assimilatory nitrate reduction (e.g., *narB/C* and *nirA*) decreased over time while denitrification (e.g., *narZ/J/I/Y/G/H*, *nirS/K*, *and nosB/C/Z*), anammox (e.g., *nirS/K* and *hzo*), and nitrification (e.g., *amoA/B*, *hao*, and *nxrB*) increased (Figure 5E). Assimilatory nitrate reduction reduces NO_3_^-^ to NH_4_^+^ for organic N synthesis. Denitrification and anammox pathways reduce NO_3_^-^ and oxidize NH_4_^+^ to N_2_, which can lead to nitrogen leaching and losses in the environment. Nitrification, or mineralization, converts reduced ammonia to inorganic nitrogen. The overall abundance pattern of N cycling-related genes leaned towards reduced assimilatory nitrate reduction and nitrogen leaching in the topsoil samples (Figure 5E). As the restoration progressed, higher levels of organic degradation and synthesis provided substrates for anammox (Figure 5E). These observations were further verified at the gene level (Figures S8F-S8K). Nitrification activity is higher in soil with a lower C: N ratio (particularly < 20:1)^67^, which is true in the SI regions (Figure S2A). To facilitate restoration and reduce nitrogen loss via nitrification, we could modify the C: N ratio by adding carbon substrates such as paper, dry leaves, and woodchips.

In the SD region, nitrogen fixation, organic nitrogen degradation and synthesis, and denitrification showed the lowest abundance (Figure S6B), as most nitrogen-fixing bacteria have symbiotic or associative relationships with plants, which are scarce in the SD region. On the other hand, the highest abundance of nitrification genes in the SD region indicates a constant loss in inorganic nitrogen.

The corroborative nature of biogeochemical cycles was evident during restoration. The increase of a more efficient HP/HB cycle for CO_2_ fixation (Figure 5A) and the negligible change in N_2_ fixation (Figure 5E) could be attributed to the limited phosphorus in the soil since the phosphorus cycle is intricately connected to carbon and nitrogen metabolism in the soil (Supplemental information; Figures S8A-S8E). Despite an observed increase in total phosphorus over time (Figure 1D), it still lagged behind the natural topsoil level^68^. The observed enhancement in methylotrophic methanogenesis and anaerobic CH_4_ oxidation (Figure S7A) indicated the presence of sufficient SO_4_^2-^, NO_x_^-^, Fe, and Mn as electron acceptors^63–65^ and high salinity^62^ in the soil. Concordantly, sulfur-reduction pathways (Supplemental information; Figure S7G), denitrification pathways (Figure 5E), Fe concentration, and EC (Figure 1D) increased over time. The increased sulfur oxidation (Figure S7G) implied that the desert soil contained adequate organic matter and a higher availability of other nutrients^69,70^, which was further supported by the rise in catabolic carbohydrate-active enzymes like GHs, CEs, and PLs (Figure S6D), as well as the enhancement of most macronutrients and micronutrients (Figure 1D). Consequently, although nutrient levels were lower than typical natural topsoil, biogeochemical cycles were systematically remodeled to ensure energy balance and nutrient accumulation in the restoring environment. Importantly, microorganisms from bacterial, archaeal, fungi, and even viral domains all contribute to the biogeochemical remodeling processes, often in a consistent manner (Figures 5 and S6-S8).

We analyzed the sources of the variation (Figure 5F) in nutrient cycles in the SD and SI regions. All five biogeochemical cycles were significantly associated with soil biophysiochemical attributes, with C-cycle (77.7%) and S-cycle (49.1%) being the highest. Among them, the Ks played an important role in all nutrient cycles except CH_4_-cycle and C-cycle.

In summary, the coordinated rewiring of biogeochemical cycles underlies the drastically changing soil environment, with microorganisms and hydraulic properties as the major contributing factors.

### Functional impact of drip irrigation and monoculture on desert soil microbiome

We were interested in how the different irrigation (Figures S9 and S10) and planting methods (Figure S10) impact the soil microbiome and metabolome landscapes. In the DI regions, the Shannon diversity of bacteria and archaea decreased over time, the opposite trend of SI regions (Figure S9A). Regression analyses (*p.adj* < 0.1) showed that the phyla involved in nutrient cycles generally decreased, except for the photosynthetic Cyanobacteria. Overall, the DI regions had higher Cyanobacteria, while the SI regions had higher Proteobacteria (Figure S9F). We did not observe significant differences in the metabolome of the DI regions.

The species interaction network from DI regions suggested a more centralized community (Figure S10A). In contrast to SI, the DI network decreased community stability over time (Figure S10B). Thus, DI and SI strategies induced opposite trends at the network level, and the SI method may benefit the microbiome diversity and stability long term.

We then analyzed the functional diversity of the soil microbiome in the DI regions. Interestingly, in contrast to the SI regions (Figure 4C), the relative abundance of photosynthesis-related pathways increased over time (Figures S10C and S10F), while the decreased pathways included carbohydrate and amino acid metabolism. Meanwhile, the increasing GTs and decreasing PLs indicate stronger assimilative functions and weaker catabolic functions (Figure S10G). Additionally, for the biogeochemical cycles, the increasing Calvin cycle and nitrogen fixation functions (Figure S10H) were probably simulated by the enhancing FVC (Figure S2B) and Cyanobacteria (Figure S9B). Overall, the DI regions showed higher anabolic and weaker catabolic functions during restoration, the opposite trend of SI regions.

At the monoculture site, the Shannon diversity of bacterial and fungal communities in the NBC group was higher than in the BC and NR groups (Figure S10I) partly because Cyanobacteria (21%) dominates in the biocrusts (Figures S10J-S10K). Actinobacteria was higher in the SD2 and NBC samples, resembling the typical desert sands. The higher abundance of photosynthetic Proteobacteria and Cyanobacteria (Figure S10K) is consistent with higher carbon in the BC and NR groups (Figure S2D). The BC and NBC groups also showed differences at the functional and metabolic levels (Figures S10I-S10O).

These results indicate that the polyculture and SI methods performed better than the monoculture and DI methods at the ecological and microbial functional levels.

### Microbes promote soil health via metabolites

To further investigate the impact of the microbiome on soil health, we trained the gradient-boosting regression models on genus and metabolite profiles to predict Ks and carbon, arguably the most important physical and chemical properties of the soil, with high accuracy (R = 0.926 and R = 0.935, respectively; Figure 5G; Methods). Actinobacteria and Proteobacteria genera dominated the top 20 variables for predicting Ks and carbon (Figures 5H and S11A). Three metabolites, including the beneficial 3-Indoleacrylic acid, were among the top 20 variables for predicting carbon. Some genera and metabolites, which were positively correlated with carbon or negatively correlated with Ks (Figures 5H and S11A), may benefit the restoration process.

We next explored the potential causal relationships between the microbiome and soil health. Among different kingdoms, only the Shannon index of fungi in the soil positively correlated with the restoration duration (Spearman test, Rho = 0.37, *P-value* = 5.649 × 10^-8^). Fungi showed higher resistance than bacteria to changes in water availability, enabled by hyphae that may traverse air-filled soil pores to access nutrients and water^11^. We used structural equation modeling (SEM) to investigate the direct and indirect impact of the restoration duration and fungal communities on macronutrients. The results showed that fungal diversity directly and positively affected the soil’s carbon, nitrogen, and phosphorus levels but did not significantly affect sulfur levels (Figures 6A, 6B, S11B, and S11C). The restoration duration had a direct, positive effect on carbon and nitrogen levels (Figures 6A and S11B) and an indirect, positive effect on phosphorus and sulfur levels through microbial biomass (Figures 6B and S11C). The irrigation methods also affected soil nutrition through microbial biomass and fungal diversity (Figures 6A, 6B, S11B, and S11C). Overall, SEM results demonstrate that fungal diversity, restoration duration, and irrigation methods may have a causal impact on soil biophysiochemical properties.

**Figure 6.**
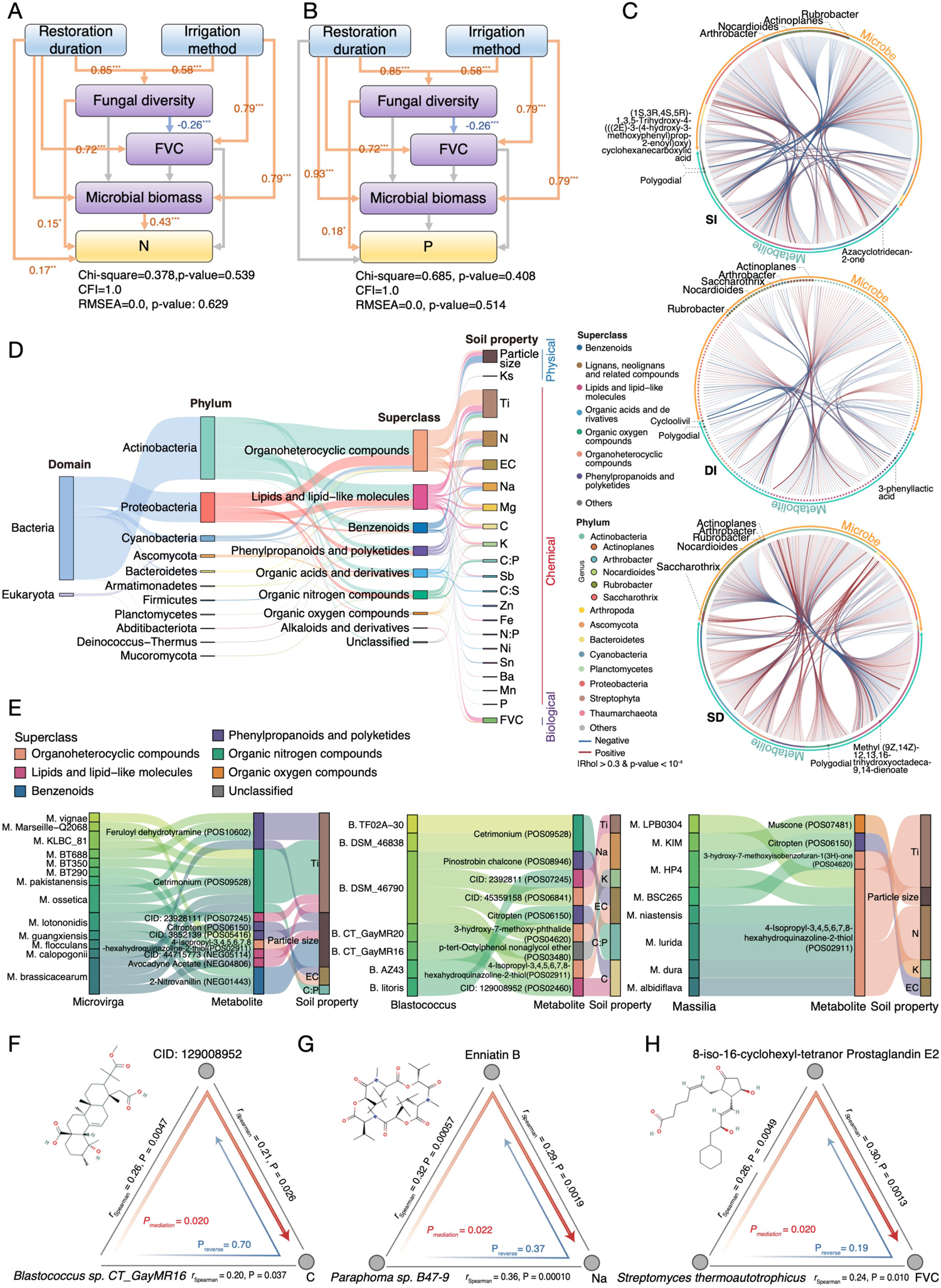
Potential causal relationships between the microbiome, metabolites, and soil health (A-B) The SEMs on nitrogen (A) and phosphorus (B). Blue lines indicate negative effects, and orange lines indicate positive effects. Grey denotes non-significance. \*\*\**P* < 0.001; \*\**P* < 0.01; \**P* < 0.05. (C) Microbes-metabolite interactions at the polyculture site. Dot represents a specie or a metabolite. The color of points represents the biological or chemical taxonomic classifications. Edge colors denote positive (red) or negative (blue) correlations. Chemicals negatively correlating with multiple species are labeled. (D) Sankey diagram showing the 240 significant mediation linkages between microbiome, metabolites, and soil health properties. The sizes of blocks indicate the number of mediation linkages. (E) Sankey diagrams showing the mediation analysis results of three representative genera. The colors of metabolites represent the superclasses. (F) Mediation effect of *Blastococcus sp. CT_GayMR16* on the carbon as mediated by CID: 129008952. (G) Mediation effect of *Paraphoma sp. B47-9* on the Na as mediated by Enniatin B. (H) Mediation effect of *Streptomyces thermoautotrophicus* on the FVC as mediated by 8-iso-16-cyclohexyl-tetranor Prostaglandin. Coefficients and *P*-values (Spearman test) for each correlation are shown in the mediation plots. Direct mediation (microbe-metabolite-soil health; red arrow) and reverse mediation (microbe-soil property-metabolite; blue arrow) are illustrated. Mediation *P*-values were calculated based on 999 bootstrap replicates. See also Figure S11.

Microbes use metabolites, such as quorum-sensing molecules and antibiotics, for synergistic and competitive purposes. We profiled the microbe-metabolite correlations in the polyculture site (Spearman, |Rho| > 0.3 and *p* < 10^-4^) and identified 365 (pos/neg: 168/197), 182 (86/96), and 384 (283/101) correlations in the SI, DI, and SD regions, respectively (Figure 6C). The SD region showed significantly more positive correlations than other regions, indicating pronounced synergistic relationships under extreme environments (Figure 6C). The same genus can interact with metabolites differently in different regions (Figure 6C). For example, the genera *Nocardioides, Saccharothrix,* and *Rubrobacter* were predominantly negatively correlated with metabolites in the SI and DI regions but positively correlated with metabolites in the SD region. *Actinoplanes* was positively correlated with various metabolites in the DI regions but exhibited predominant negative correlations in the SI regions, while *Arthrobacter* showed an opposite trend. Last but not least, many known and potential antibiotic chemicals were negatively related to diverse microbes (Figure 6C).

To evaluate whether metabolites can mediate the role of microbes on soil health, we applied bi-directional mediation analysis on the SI samples, revealing 240 mediation linkages (adjusted-P_mediation_ < 0.05 and adjusted-P_inverse mediation_ > 0.05; Method, Figure 6D).

Most of these linkages were related to the microbial impact on soil particle size, Ti, N, EC, Na, and Mg (Figure 6D). Actinobacteria, Proteobacteria, Cyanobacteria, and Ascomycota (Fungi) were commonly involved (Figure 6D). The top 5 genera ranked by the number of linkages were *Arthrobacter, Microvirga, Nocardioides, Blastococcus, and Massilia,* each affecting soil properties through different metabolites (Figures 6E, S11D, and S11E). For example, the nitrogen producer *Massilia*^71^ may impact soil nitrogen via lipids and lipid-like metabolites. The major components of biocrust, *Blastococcus* and *Geodermatopliu,* showed diverse mediation linkages of soil carbon (Figures 6F and S11E). Numerous species, primarily from the genera *Agromyces*, *Microvirga*, and *Mycolicibacterium*, produce metabolites that mediate the decrease in particle size (Figure S11F). We also found that enniatin B, a fungal metabolite that forms form complexes with alkali metal ions^72^, could mediate the impact of *Paraphoma* sp. B47-9 on Na concentration (Figure 6G). Another interesting example is the mediation of 8-iso-16-cyclohexyl-tetranor Prostaglandin E2 (8-iso PGE2) by *Streptomyces thermoautotrophicus* on FVC (Figure 6H). Prostaglandins (PGs) have been detected in many plants and certain microorganisms—different PGs impact plant growth differently, including accelerating flower formation^73^.

The combined statistical learning, correlational, and causal analyses identified microorganisms and metabolites as predictors and potential biotic agents for desert restoration.

### Evolutionary adaptations in the desert soil microbiomes

The microbiome is constantly evolving. The deep-sequencing data enabled us to examine intraspecies variation. Using the uniquely mapped sequencing reads, we investigated the genomic evolutionary landscapes of the top 100 species at the polyculture site. We calculated the single-nucleotide polymorphism (SNP) density and nucleotide diversity (π) across all filtered genomic positions across species from different domains of life (Figure 7A; Methods). We identified 14.37M SNPs in the selected 100 pan-domain species. As expected, we found that SNP density was highly concordant with nucleotide diversity (R = 0.95, *p* < 2.2e-16) (Figure S11G), and there was greater genomic diversity in species with higher coverage across all domains (R = 0.82, *p* < 2.2e-16; Figure S11H).

**Figure 7.**
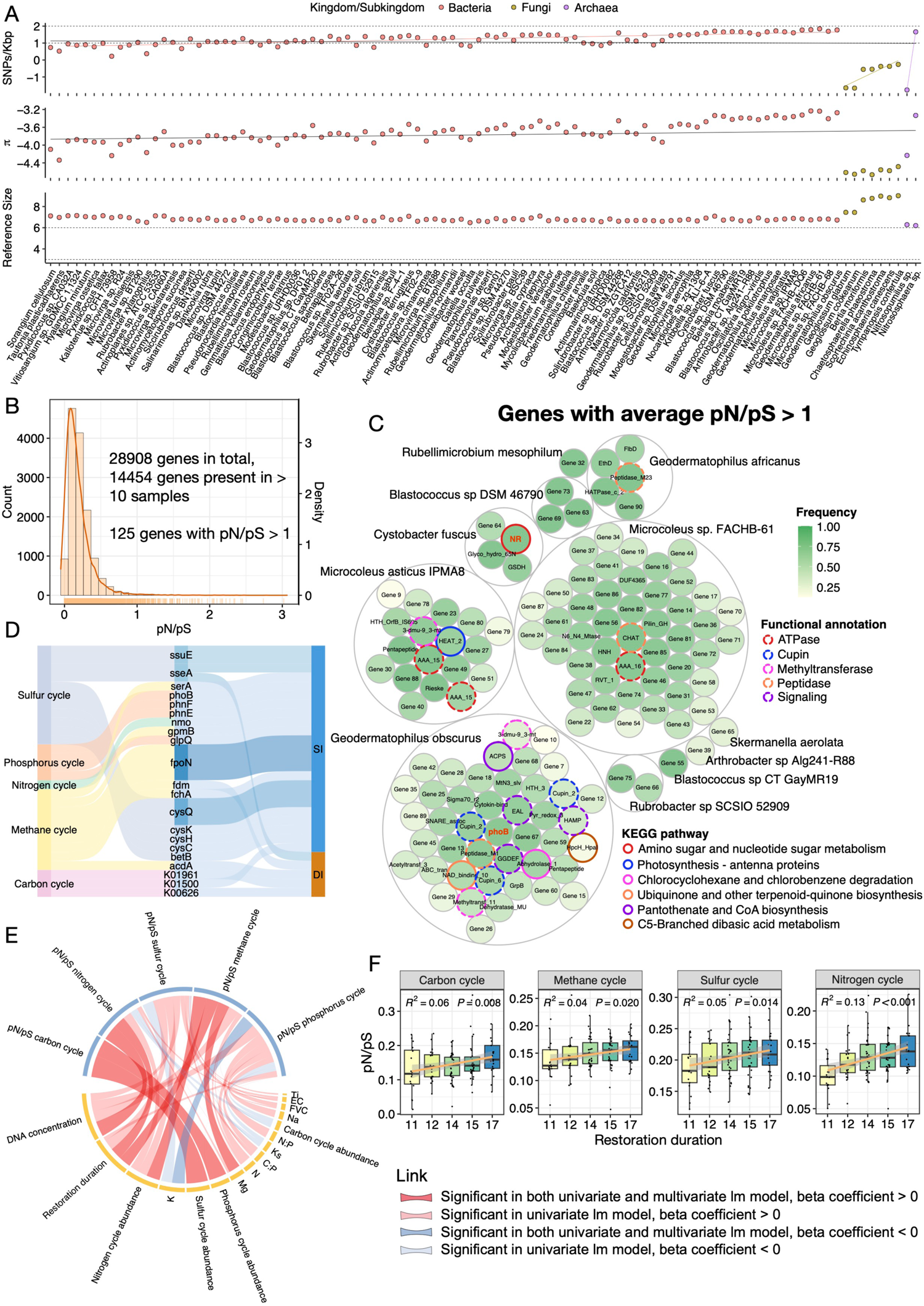
Evolutionary adaptations of desert soil microbes during restoration. (A) Population genetics analyses of the top 100 species from bacteria, fungi, and archaea. First row, SNP density, the two dashed lines denote 100 and 10 SNPs/Kbp, respectively; The colored linear regression lines indicate that SNP density positively correlated with coverage in bacteria (red), fungi (brown), and archaea (purple); the second row, nucleotide diversity (π or pi); and the third row, reference genome size, dashed line denotes 10^6^ bps. All values are in the log scale (base 10). (B) The distribution of pN/pS of all genes for the top 30 abundant species (see Methods). Histogram (left y-axis) and density plot (right y-axis) were shown. (C) The genes with pN/pS greater than one and present in at least ten samples. Each small circle represents a gene filled based on the frequency of occurrence in samples. 125 genes from 11 species are shown. A larger gray circle-packed gene of the same species. Genes were annotated by Pfam (gene name; black text) and KEGG pathways (colored circles) and marked by functional annotation (colored dashed circles). Unannotated genes were represented by “Gene number”. (D) The biogeochemical cycling genes that experienced positive selection. The three columns from left to right refer to biogeochemical cycles, biogeochemical cycling genes, and irrigation methods. The more samples with pN/pS > 1, The darker-colored links indicate more samples with pN/pS >1. (E) The relationship between the pN/pS of the five biogeochemical cycles (top; blue) and soil biophysiochemical properties (bottom; yellow). Both univariate and multivariate models were shown. The multivariate models incorporated all variables at the bottom. The more significant the association, the thicker the link. (F) The significant temporal patterns of pN/pS of the biogeochemical cycles in SI regions. The R-squares and p-values were calculated by the linear model regression.

We calculated the pN/pS of all genes with sufficient coverage (Methods). Most of the genes had pN/pS close to 0, which is expected as mutations are generally deleterious. However, we observed more than 125 genes with positive selection signatures (Figure 7C). They were related to ATPase, Cupin, different types of transferases, peptidase, signaling transduction, pili, and flagellar (Figure 7C). Two biogeochemical cycling genes, NR and phoB, also experienced positive selection. In contrast, genes with pN/pS close to zero were subjected to purifying selections, and they are mostly involved in translational, transcriptional, and essential functions (Figure S11I). We focused on genes participating in biogeochemical cycles, and numerous genes in different steps of nutrient metabolisms were identified, especially in the sulfur and methane cycles (Figure 7D). This is surprising as genes in biogeochemical cycles are normally considered constitutive in cellular organisms, which should be subjected to purifying selections. The relatively fast-changing restoring environment likely drives the positive selection of biogeochemical genes.

We further analyzed how biophysicochemical factors influence the pN/pS of the major biogeochemical pathways at the polyculture site (Figure 7E). DNA concentration, restoration duration, N, S, P cycling gene abundance, K, and Mg were positively correlated with the pN/pS of different biogeochemical pathways in both uni- and multivariate regression models. Interestingly, the abundances of different cycles did not directly correspond to their own pN/pS ratios. For example, the pN/pS of the C cycle was positively correlated with the relative abundance of the S and P cycles. We further examined how restoration duration impacts the evolution of biogeochemical cycles, and all cycles except for P showed significant correlations with restoration duration, also validated at the gene level (Figures 7F and S11J).

In conclusion, the microbiome evolutionary analyses indicate that the desert microbiome underwent rapid evolutionary changes related to biogeochemical remodeling, among other enzymatic and motile functions.

## Discussion

Ecological restoration of damaged lands is one of the top priorities in the Sustainable Development Goals (SDGs) proposed by the UN. Restoration of the desert landscape has proven to be especially difficult. Vegetation restoration has been implemented and proven to be efficacious in facilitating the restoration process^7–9^, but few studies have explored the microbial role during long-term desert restoration, despite its extensive interactions with plants and soil.

We used space-for-time substitution to investigate the long-term restoration process of the desert ecosystem with deeply sequenced metagenomic, metabolomic, and soil physicochemical data. Most desert-related studies used marker genes such as 16S/18S rDNA/rRNA^25^, providing limited functional and evolutionary information. The deep-sequencing data provided the necessary genomic information to construct a redox-based biogeochemical model for desert ecosystems. The multi-dimensional analyses depicted detailed physicochemical, compositional, functional, biogeochemical cycling, ecological, and evolutionary landscapes during the long-term desert restoration, which would appeal to colleagues in the microbiome, microbiology, ecology, evolution, soil science, bioprospecting, antibiotics discovery, etc.

Our results also provided a rich context of existing desert microbiome studies. We showed that sand dune microbiomes exhibit distinct characteristics, with a higher relative abundance of CO_2_ fixation, methanogens, and methanotrophs; and a lower relative abundance in nitrogen, phosphorus, and sulfur cycles. Many metabolites positively correlated with microbial abundance, especially in the SD region, indicating synergistic relationships in extreme environments (Figure 6C). In contrast, numerous metabolites inversely correlated with microbes In the SI regions, indicating the potential antimicrobial activities (Figure 6C). Importantly, these characteristics underwent extensive changes during restoration, suggesting that the desert microbiome can be extensively remodeled *in situ* to facilitate restoration.

Restoring damaged ecosystems requires at least two physicochemical improvements: hydraulics and nutrient accumulation. Causal and predictive analyses have highlighted the potential role of microbes and metabolites in promoting soil health (Figures 5G and H; Figure 6). For instance, *Agromyces* and *Mycolicibacterium* have been identified as producers of bio-erosive compounds that reduce soil particle size, thus enhancing hydraulic properties (Figure S11F). Additionally, the introduction of *Caulobacter*, which is commonly found in endospheres or rhizospheres, along with 3-Indoleacrylic acid or *Aspergillus*, a filamentous fungal genus, may contribute to carbon accumulation (Figure 5H).

Through comprehensive functional analyses spanning all domains of life, we demonstrated that bacteria, archaea, fungi, and even viruses play important roles in biogeochemical remodeling. Insights from biogeochemical remodeling have strong potential for field applications. For example, adding phosphorus-supplying organic materials like manure or wood ash can alleviate carbon and nitrogen fixation restrictions resulting from phosphorus deficiency (Figure S8A). Similarly, incorporating pure carbon substrates such as dry leaves or woodchips to improve the carbon-to-nitrogen ratio can reduce nitrogen loss and facilitate nitrogen fixation (Figure 5E). Furthermore, introducing microorganisms capable of assimilatory nitrate reduction, such as members of Alphaproteobacteria, Planctomycetia, Actinomycetia, and Bryopsida, can further enhance nitrogen fixation (Figure S8F). Notably, members of Actinomycetia, Betaproteobacteria, Chloroflexia, and Cytophagia (Figure 5C) utilize HP/HB cycle to fix CO_2_, which holds promise for metabolic engineering because of its efficiency, thermostability, and rapid metabolic pathway kinetics in either aerobic or anaerobic environments. Alternatively, engineering the HP/HB cycle into other desert extremophiles may further facilitate restoring damaged lands.

Our study has several potential limitations: (1) The organismal sequence database needs to be expanded. We do not have genomic data on the planted species, and 58.96% of DNA reads could not be classified. (2) The chemical database needs to be expanded. Of the 18885 putative compounds, only 1449 had level-2 annotations. Further research is needed to confirm the numerous predicted features. (3) Studies in more similar experimental restoration sites can further confirm the findings.

Studies on the restoration of ex-mined areas^74^ and soil after a wildfire^75^ have demonstrated that soil microbiomes may facilitate repairing damaged ecosystems. However, the detailed functional role of the microbiome in different types of damaged ecosystems could vary significantly. For instance, a study on the impact of vegetation restoration in ex-mined areas showed that genes involved in transposable elements, virulence, defense, and stress response increased during restoration. Surprisingly, microbiome taxonomic and functional changes suggested a shift from a copiotrophic to an oligotrophic state, and the key N-cycle-related genes did not change^74^. In a post-wildfire soil microbiome study, the severity of the fire damage was directly correlated with the bacterial gene abundances for heat resistance, fast growth, and pyrogenic carbon utilization, which might enhance post-fire survival^75^. Systematically expanding our knowledge of the functional and ecological role of the microbiome in natural or active restorations of damaged ecosystems is essential in developing customized effective microbial restoration approaches, which may eventually have applications beyond Earth. For example, terraforming Mars and extra-terrestrial agriculture would require the restoration of desert-like Martian soil.

## Acknowledgments

This study was supported by the Zhejiang University LSI start-up funding. We greatly appreciate Alxa Tengger Desert Lockside Ecological Public Welfare Project Base, the Alipay Ant Forest-Society of Entrepreneurs & Ecology, and Mr. P. Li for their help in sampling. We thank Mr. J. Jiang for supplying aerial photographs and Y. Chen, C. He, K. Yuan, and K. Tang for their help in the analyses.

## Author contributions

C.J. and Q.C. conceived the study. C.J., Q.C., L.J., and Z.L. collected all samples. Q.C., M.Y., and Z.L. performed experiments. Q.C. led the bioinformatic analyses. X.W. performed taxonomy classification, co-assembly, and evolutionary analyses. M.Y. and Z.H. analyzed metabolomic data. L.J. performed SEM and mediation analyses. Z.L. annotated the CRISPR Array. C.P. performed network analysis. D.T. estimated the FVC. X. Wu and J. S. contributed to the sampling. M.J., L.H., Q.L., X.Z., P.G., and C.Y. contributed to the analyses. Q.C. and C.J. drafted and revised the manuscript with input from other authors.

## Data availability

The raw sequencing reads were submitted to ENA under project ID PRJEB60065.

## Code availability

The main analysis script was written in the Rmarkdown format and deposited on GitHub at https://github.com/Q-chen27/Desert_ecosystem_restoration.git.

## STAR Methods

### Sample collection

We implemented the space-for-time substitution (SFT) method and standardized protocols^76^ to collect samples from two restoration sites located in the Alxa Left Banner, near Bayanhot, on the eastern edge of the Tengger Desert in Inner Mongolia, China (Figure S1A). The desert is expanding in size. This region is a temperate desert arid climate (Köppen system) characterized by heavy sandstorms, extremely low precipitation, little cloud cover, and strong evaporation. This area also features considerable seasonal temperature variation, periods of drought, and severe desertification. The annual average temperature is 7.2 degrees Celsius, the mean sunshine time is 3316 hours, the frost-free period is 120 to 180 days, and the annual precipitation is 71.44–116.60 mm, with 2900–3300 mm annual evaporation. The annual average wind velocity is 2.48–2.79 m/s^77^. The soil at the site is aeolian sandy soil.

From October 13 to 15, 2020, we collected 1490 samples (pooled into 298 testing samples) at the polyculture site. Briefly, 10 grams of topsoil (depth: 0 to 5.0 cm) were collected at each of the five sites within an approximately 1 m^2^ area in a diagonal cross pattern and pooled together to store in sterile 50 ml polypropylene Falcon tubes (Corning, USA) aseptically mixed as a sample. Table S1 shows detailed sampling information. The entire restoration site was fenced. Each area was planted with similar proportions of *Hedysarum scoparium, Ammodendron, Caragana korshinskii, and Calligonum mongolicum*. Lines represent the location of the sampling sites in the region (Figures 1A and S1A). For each sampling lane, 10 to 20 samples were collected, spaced 15 meters apart. The samples were named based on the restoration duration. Samples collected along the fence, covering the 2003 to 2006 restoration area, were defined as the Inside Fence (IsF) group. Samples collected near but outside the fence towards the Tengger desert were characterized as the OsF group. *Agriophyllum squarrosum* was present in the OsF group, where camels were periodically grazing. Pure sands collected from sand dunes (SD) were defined as a negative control group because there was few biotics. Importantly, the biocrusts would naturally form at the polyculture site, but the site workers will disrupt biocrust formation to facilitate vegetation restoration as their common practice.

The monoculture site was also located in the Alxa Left Banner, to the north of the polyculture site (Figure S1A), which is funded by the SEE Foundation. On October 15, 2020, we collected 420 samples (pooled into 84 testing samples) with the same method described above. Samples were equally distributed in every sampling area. This site is characterized by the monoculture method, with only one species mentioned previously planted in each region. Only manual subsurface irrigation is used. For each restoration region, 10 samples were collected from non-biocrust areas, and 4 samples were collected from biocrust areas. Biocrust is a craggy, often dark or burnt-looking carpet stretching between shrubs and grasses in arid lands. It is a community of lichens, bryophytes, and Cyanobacteria that live on the soil surface of drylands and are vulnerable to soil disturbance^78^. The natural recovering (NR) region was formed due to the restoration region blocking the movement of sands. We also collected nearby sand dunes (SD2) as a control group.

All the 382 testing samples were aliquoted to three tubes after vortex on a Vortex Genie II vortex mixer (Scientific Industries Inc., Bohemia, NY) on the sampling day and stored at −20°C. The samples were soon transported on dry ice to Zhejiang University to freeze at −80°C for further experiments (received on October 18, 2020). Detailed metadata information can be found in Table S1.

### FVC estimation

Fractional vegetation cover (FVC) is defined as the projected percentage of the total vegetated area (leaves and stems)^79^, reflecting the size of the plant’s photosynthetic area and the density of vegetation growth. The Normalized Difference Vegetation Index (NDVI) is widely used for monitoring vegetation-covering areas on a remote sensing setup^80^. The Landsat8OLI-TRIS satellite digital data with a 30 m * 30 m resolution of the two restoration sites in this study were downloaded from the Geospatial Data Cloud (http://www.gscloud.cn). We acquired the satellite images captured on August 18, 2019.

The average cloud cover was 0.95. We used the ENVI software to pre-process the satellite image, which includes two steps: radiometric calibration and atmospheric correction. The preprocessed images were further used to extract NDVI and estimate FVC.

The NDVI assesses vegetation greenness and is useful in understanding vegetation density; the formula used for calculation is: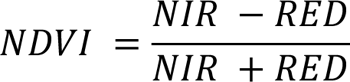

Where NIR is near-infrared reflectance, and RED is visible red reflectance. The NDVI is calculated as the difference between the reflection value of the near-infrared band and that of the red band, divided by the sum of these two values^81^.

FVC is calculated based on the pixel dichotomy model^82^. The model assumes the pixel is only composed of vegetation and non-vegetation cover parts, and the spectral information is only linearly synthesized by these two parts, with the proportion of their respective areas in the pixel set as the weight of each part. The formula is: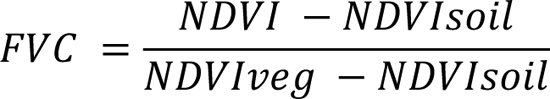
where NDVI is the weighted average of vegetation and non-vegetation regions; NDVIsoil is the vegetation index of the bare soil pixels, and NDVIveg is the vegetation index of the vegetation pixels. The 95% cumulative percentage of NDVI based on the global data is the value of NDVIveg. NDVImin and NDVImax are obtained at 5% and 95%, where NDVIsoil = NDVImin and NDVIveg = NDVImax are substituted into the formula to calculate the FVC. After calculations, FVC is extracted according to the GPS coordinates of each actual sampling point and the number of samples.

### Soil size, texture, and hydraulic conductivity analysis

We used dynamic image analysis with a Camsizer X2 (Retsch Technology GmbH, Newtown, PA) to characterize the size distribution of all samples. We pooled 2 to 4 samples from the same sampling lane (DI and SI regions) of the same sampling region. The Camsizer X2 has a measurement range of 0.8 μm – 8 mm. The wide range of analysis allows for efficient and consistent quantification of soil size parameters without careful sieving (about 2 minutes per sample and 5 g of sample per 100,000 particles). Three key size parameters measured were equivalent circle diameter (xarea), minimum chord diameter (xcmin), and maximum Feret diameter (XFemax). Data can be output as volume fractions or particle number distributions (PND; the number of particles in each size fraction)^83^. Soil texture, defined by the relative percentage of sand (2.0 – 0.05 mm), silt (0.05 – 0.002 mm), and clay (less than 0.002 mm) particles, is an important soil environmental variable^84^. We used PND to calculate soil texture and hydraulic conductivity, a vital parameter for predicting water movement and dissolved contaminants through the soil ^85^. The following formula was used^85^: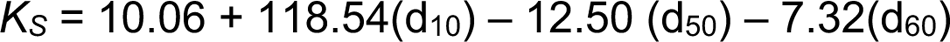

Where d_10_, d_50_, and d_60_ are the soil particle diameters (mm) at which 10%, 50%, and 60% of all soil particles are finer (smaller) by weight, and Ks is the saturated hydraulic conductivity expressed in m/day. d_10_, d_50_, and d_60_ are calculated using PND, assuming that each particle has the shape of a circle with the same density. The higher the Ks, the lower the hydraulic conductivity.

### Evaluation of soil physicochemical properties

Physicochemical analyses were performed using standard ISO/IEC 17025 procedures. Physicochemical parameters analyzed included: moisture, pH, electrical conductivity, macronutrients (C, N, P, S, K, Mg, and Ca), micronutrients (Fe, Mn, Cu, Zn, Mo, Ni, Na, and Co), and other elements (Ti, Al, As, Ba, Cr, Cd, Pb, Sb, and Sn). Soil moisture was determined by oven-drying at 105°C to a constant weight (Shanghai YIHENG Technical Co., Ltd., Shanghai, China) and expressed as a percentage of soil water to dry soil weight. We measured soil pH and conductivity from soil water suspension (1:5 v:v) with a digital pH meter (Mettler–Toledo Instruments Co. Ltd., Shanghai, China). We prepared samples to analyze macronutrients, micronutrients, and other elements by grinding them and sieving them through a 0.15 mm mesh. We measured the total C and N of all samples using a Vario Max CN Analyzer (Elementar Analysensystem GmbH, Hanau, Germany). Other nutrients and heavy metals of the DI, SI, and SD from the polyculture site were extracted with pure acid (6 mL HNO_3_ + 2 mL HCl + 1 mL HF; Analytical Reagent; China National Pharmaceutical Group Corporation, Beijing, China) and measured using inductively coupled plasma atomic emission spectroscopy (ICP-AES, JAC IRIS/AP, Thermo Jarrell Ash Corporation, Franklin, USA).

### DNA extraction and library preparation

We extracted total DNA from 500 to 700 mg of each testing sample using the Power Soil DNA Isolation kit (QIAGEN, Hilden, Germany) with minor modifications. DNA concentration was measured by Qubit Fluorometric Quantitation (Thermo Fisher Scientific, Waltham, USA). DNA libraries were prepared using the VAHTS Universal Plus DNA Library Prep Kit for Illumina® (Vazyme #ND617, Nanjing, China) with adaptors and primers from the VAHTSTM Multiplex Oligos set 5 for Illumina® (Vazyme #N322) as described by the manufacturer. DNA input for library preparation was 4 ng except for the SD and SD2 controls, which used 500 pg due to extremely low DNA yield. We assessed the size and quality of libraries on a Qsep100 (BiOptic, New Taipei City, China). We sequenced the libraries using the Illumina NovaSeq 6000 platform (2 × 150 bp) (Illumina Inc., San Diego, CA) run by Novogene Co., Ltd (Beijing, China).

### Contamination considerations

Because our experimental protocol was designed to extract and prepare libraries from sand dune samples, we carefully evaluated the potential for contamination using the following approaches:

1. The recommended sample input (250 mg, Power Soil DNA Isolation kit) did not yield sufficient DNA for desert soil samples in the preliminary experiments; we increased the input to 500 mg for samples collected from restoration sites. 700 mg was used for samples collected from negative sand dune (SD) controls;
2. We included two types of blank controls to evaluate potential contamination from the experimental processes, including the Power Soil DNA Isolation kit and the library preparation kit. 1. Blank samples (RNase-free and DNases-free water) were extracted with the Power Soil DNA Isolation kit and then used for library preparation. 2. RNase-free and DNase-free water were directly used for library preparation. We could not quantify the DNA concentration of libraries after library preparation procedures with the same number of amplification cycles as the testing samples, indicating that background contamination from the kits and experimental processes was negligible;
3. Three operators performed DNA extraction and library preparation. The fact that batch effects had a very small role in our variation partition analysis suggests that batch effects are well controlled.

Together, these results indicate that contamination is insignificant in our dataset.

### Taxonomic classification of metagenomic reads

To explore the pan-domain species composition of the desert microbiome, we performed taxonomic classification for metagenomic sequencing reads with Kraken2^33^ and a comprehensive custom database of bacterial (370,424), fungal (7485), viral (41,963), archaeal (6269), viridiplantae (1612), vertebrate (1242), invertebrate (1120), and protozoal (994) genomes from the union of NCBI RefSeq and GenBank databases (numbers in parentheses indicate the number of genomes). The size of the resulting Kraken2 pre-built index is more than 3TB. Although running Kraken2 with such a large index was time-consuming, our custom database annotated significantly more reads than Kraken2’s standard database. The reference genomes were downloaded from RefSeq and GenBank on December 30, 2020.

### MAG construction and annotation

#### 1. Binning

The obtained high-quality clean reads were assembled de novo by MEGAHIT (v1.2.8)^86^ to individually generate contigs. Open reading frames (ORFs) in contigs were predicted using the MetaGeneMark (v3.38)^87^. DAS Tool (v1.1.4)^88^ was used to select the best bacterial and archaeal bins from the combination of three programs — MetaBAT2 (v2:2.15)^89^, VAMB (v3.0.7)^90^, and SemiBin (v1.0.0)^91^, resulting 2787 refined bins. Specifically, Nissen *et al.* summarized three binning modes: the single-sample approach, where each sample is binned independently; the multi-sample approach on a coassembly; or a multi-split approach^90^. Because of the size of our desert metagenomes, the multi-sample approach, which needs co-assembly with all reads, is time- and resource-consuming. Instead, we performed the single-sample approach with MetaBAT2 and SemiBin, while using the ‘binsplitting’ mode of VAMB as recommended by the developers (https://github.com/RasmussenLab/vamb). MetaBAT2 was run using its default parameters. SemiBin used a semi-supervised siamese neural network dealing with the must-link and cannot-link constraints. We used the prebuilt model of SemiBin with the command “SemiBin single_easy_bin -m 1000 --environment soil”. VAMB implemented a multi-split approach as recommended by the developers (https://github.com/RasmussenLab/vamb) and was run using the command “vamb -- minfasta 100000 -jgi depth.txt”.

#### 2. Dereplication and species-level clustering

To dereplicate the genome bins across all samples after bin refinement by DAS Tool^88^, dRep (v3.0.0)^92^ was used on the 2787 bins by the two-step clustering with the command ‘dRep dereplicate -comp 0 -con 10’. First, MAGs were divided into primary clusters using Mash at a 90% ANI. Then, each primary cluster was used to form secondary clusters at the threshold of 99% ANI with at least 10% overlap between genomes. The CheckM (v1.1.3)^93^ ‘lineage_wf’ pipeline was used to determine the completeness and contamination of the bins through the identification and quantification of single-copy marker genes. According to the MIMAG^94^ standards set up by the Genomic Standards Consortium, 1565 nonredundant MAGs were divided into high-quality MAGs (>90% completeness and <5% contamination), medium-quality MAGs (>=50% completeness and <5% contamination), and medium-quality MAGs (<50% complete with <10% contamination).

The 1565 nonredundant MAGs were further clustered into species-level genome bins (SGBs) at the threshold of 95% ANI using ‘dRep dereplicate -comp 0 -con 10 -sa 0.95’^95^. SGBs containing at least one reference genome (or metagenome-assembled genome) in the Genome Taxonomy Database (GTDB, https://gtdb.ecogenomic.org/) were considered known SGBs. And SGBs without reference genomes were considered unknown SGBs (uSGBs)^96^.

#### 3. Relative abundance

To calculate the relative abundance of MAGs, reads from each sample were mapped to the set of nonredundant MAGs using CoverM (https://github.com/wwood/CoverM).

#### 4. Taxonomic classification

Phylogenetic inference was conducted by GTDB-Tk (v2.0.0)^97^ to classify the nonredundant MAGs based on GTDB (version r207). To resolve the taxonomic distribution of the desert MAGs, phylogenetic trees were inferred from concatenated sets of single-copy marker genes (120 bacterial or 53 archaeal genes). The recovered nonredundant MAGs spanned 16 phyla, including Bacteria (1418) belonging to *Actinobacteria* (392 genomes), *Cyanobacteria* (335), *Acidobacteriota* (275), and *Proteobacteria* (116), etc., and Archaea from the *Thermoproteota* (142) and *Thermoplasmatota* (5).

#### 5. Phylogenetic analysis

Maximum-likelihood trees were generated de novo using the protein sequence alignments produced by GTDB-Tk: we used IQ-TREE (v2.0.3)^98^ to build a phylogenetic tree of the 1418 bacterial and 147 archaeal species. The best-fit model was automatically selected by ‘ModelFinder’ based on the Bayesian information criterion (BIC) score. The LG+F+R10 model was chosen for building the bacterial tree, while the LG+F+R4 model was used for the archaeal phylogeny. Trees were visualized and annotated with ggtreeExtra (v1.6.0)^99^.

#### 6. Gene annotation

Gene calling and annotation were undertaken with Prokka (v1.14.5)^100^. These MAGs were annotated as Archaea or Bacteria based on an inferred domain derived from the abovementioned genome tree.

### Metabolomics analysis

#### 1. Sample preparation

We conducted preliminary investigations based on previous studies^101–103^ to optimize experimental variables for soil metabolite extraction. LC/MS-grade isopropanol (CAS No. 67-63-0) and LC/MS-grade methanol (CAS No. 67-56-1) were from Thermo Fisher (Carlsbad, CA, USA). LC/MS-grade water (CAS No. 7732-18-5) is from Merck (Darmstadt, Germany). We performed extractions using isopropanol/methanol/water (3:3:2 v/v/v), followed by shaking on a refrigerated orbital shaker at 200 rpm for 1 h at 4°C to ensure adequate mixing and extraction. Samples were centrifuged at 3220 ×g for 15 min at 4°C, and the extracts were frozen at −80°C for 2 h. The extracts were then dried using a vacuum concentrator (ScanSpeed 40, Labogene, Denmark), and the dried residues were resuspended in 200 uL 50% methanol and vortexed. The samples were then filtered through 0.22 um centrifugal membranes and were ready for analysis.

#### 2. Sample analysis

Metabolomics analysis was performed by the ultra-high performance liquid chromatography coupled Orbitrap mass spectrometer (Thermo, Q Exactive™ HF, United States). A Hypesil Gold C18 column (100×2.1 mm, 1.9 μm) was applied and the column temperature was set at 40°C. The sample chamber was maintained at 4°C. The binary mobile phase consisted of 0.1% formic acid (mobile phase A, CAS No. 64-18-6, Thermo Fisher) and methanol (mobile phase B) for the positive mode; and 5mM ammonium acid (CAS No. 631-61-8, Thermo Fisher), pH 9.0 (mobile phase A), and methanol (mobile phase B) for the negative mode. The flow rate was set at 0.2ml/min, and the gradient was 0-1.5 minutes at 98% A, 1.5-12 minutes at 98% to 0% A, 12-14 min at 0% A, 14-14.1 min at 0% to 98% A, 14.1-17 minutes at 98% A. The mass spectrum acquisition was scanned in the m/z range 100-1500, and ESI source settings were as follows: Spray Voltage:3.2kV; Sheath gas flow rate: 40arb; Aux Gasflow rate: 10arb; Capillary Temp: 320°C. The instrument was operated in positive and negative modes, and MS/MS secondary scans were data-dependent. The QC sample was prepared by pooling aliquots of all subject samples and injected every 10 samples.

#### 3. Data processing

The mass spectrometry data were centroided and converted from the proprietary format (.raw) to the m/z extensible markup language format (.mzML) using ProteoWizard (MSConvert tool, ver. 3.0.21094)^104^. After comparing the upstream processing parameters optimization tools: IPO^105^, Autotuner^106^, and SLAW^107^, we chose SLAW for parameters optimization, mass feature extraction, isotopic patterns extraction, peak grouping, and deisotope. Finally, the feature quantification table results (.CSV) and MS^2^ consensus spectra (.MGF) were exported for subsequent analysis.

#### 4. Data annotation

Metabolite annotation was performed using a two-step approach. First, metabolites were identified based on accurate mass (± 0.01 Da) and MS^2^ similarity against public databases. Specifically, for MS^2^ reference database construction, we integrated open-source MS^2^ databases from the GNPS community (https://gnps-external.ucsd.edu/gnpslibrary, 495,662 spectra, from 2022-6-18), MoNA dataset (https://mona.fiehnlab.ucdavis.edu/, 1,951,233 spectra, from 2023-2-7), as well as the NIST2020 Tandem Mass Spectral Library (1,143,815 spectra). Then MS^2^ consensus spectra (.MGF files) were annotated with the NIST MSPepSearch software (https://www.nist.gov/) using the dot-product algorithm, and the cutoff was set as 600. Besides, MS^2^ consensus spectra files were also uploaded to GNPS (http://gnps.ucsd.edu) and analyzed with molecular library search workflow using a modified cosine algorithm for supplementary annotations^108^. According to Metabolomics Standards Initiative (MSI)^109^, all metabolites annotated by spectral match were considered level 2 annotation. In addition, the features with MS^2^ spectra that were not matched in public databases were analyzed by SIRIUS/CSI: FingerID^110,111^ to acquire an MSI level 3 annotation. Most of the analyses focused on metabolites with level 2 annotation.

#### 5. Tree construction

The annotated and predicted compounds were deduplicated based on the name and InChIKey, resulting in 18,885 unique metabolites. The chemical classification information was extracted using InChIKeys to create a hierarchical classification table. This table was then converted into a tree structure using Graphlan^112^ and the ape^113^ package in R and uploaded to iTOL^114^ for visualization.

#### 6. Network analysis of metabolites

We performed NetID network analysis on all the obtained MS^2^ consensus spectra. The links between spectra were evaluated based on mass differences reflecting relationships such as adjuncts, fragments, isotopes, or biochemical transformations, resulting in a relationship network between annotated and unannotated features. The network structure of NetID was derived and input into Gephi 0.10 to visualize.

#### 7. Pathway analysis

We converted 1,449 annotated compounds into PubChem CID, which are compatible with HMDB, KEGG, Reactome, and WikiPathways using a web-based database (http://cts.fiehnlab.ucdavis.edu/batch). We used c-means clustering and GAM to identify dynamically changing metabolites and performed pathway enrichment using the RaMP (2.0 API R version)^115^. This API includes pathways and metabolites from HMDB, KEGG, Lipid-MAPS, WikiPathways, Reactome, and CheBI.

### General statistical analysis and data visualization

The primary statistical analyses were performed using customized python (2.7.18) and shell scripts (Linux) and Rstudio and R (4.1.2), with visualizations in Rstudio and R. Most packages used were downloaded from the Bioconductor project (https://www.bioconductor.org/). The main packages used include abind (1.4-5), ade4 (1.7-18), adephylo (1.1-11), adespatial (0.3-14), AEM (0.6), ape (5.6-1), beeswarm (0.4.0), BioManager (1.30.16), devtools (2.4.3), dplyr (1.0.8), Cluster (2.1.2), factoextra (1.0.7), ggbeeswarm (0.6.0), ggplot2 (3.3.5), ggpubr (0.4.0), ggsci (2.9), ggrepel (0.9.1), ggtreeExtra (1.4.1), glmnet (4.1-4), reshape2 (1.4.4), RColorBrewer (1.1-2), tidyverse (1.3.1), vegan (2.5.7), gam (1.22-1), and viridis (0.6.2). The source code for the R analyses is available in the supplementary code.

We tested differences between univariate variables such as taxonomic and functional diversity using non-parametric statistical tests (Wilcoxon test, Kruskal-Wallis, and Spearman correlation) unless the data were normally distributed. Data were analyzed using a generalized additive model (GAM, the smooth class was invoked directly by *s* terms with default *df*) and a generalized linear model (*link = “log”*) with the Benjamini-Hochberg method to control for False Discovery Rate (FDR < 0.05). Taxonomy with relative abundances below 0.001% was removed^116^. The Counts Per Million (CPM) method was used for inter-sample normalization, followed by hyperbolic arcsine (arcsinh) transformation (asinh function in R) for the within-sample normalization^23,117^. Statistical analyses and visualizations were conducted with arcsinh-transformed CPM values (aCPM) unless otherwise noted, such as CAZymes annotation. Annotated phylogenetic trees were generated using customized python scripts and the R package ggtreeExtra. The alpha diversity (within-sample diversity) summarizes the structure of an ecological community in terms of its richness (number of taxonomic groups), evenness (distribution of abundances of the groups), or both^118^. Here, we used the Shannon Index to measure alpha diversity with the following formula at the species level: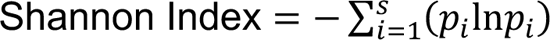

Where:

Σ: A Greek symbol that means “sum” ln: Natural log

pi: The proportion of the entire community made up of species i

The higher the value of the Shannon Index indicates the higher the diversity of species in a particular community.

Principal Coordinates Analysis (PCoA) based on the Bray-Curtis distance at the species level was used instead of correspondence analysis (CA) to discriminate the profiles of samples originating from different duration using the vegan package. This decision was based on the observation that the environmental gradient axis was shorter than 2 in the detrended correspondence analysis (DCA)^119^, indicating that the environmental gradients in our dataset are rather short and linear, making PCoA analysis feasible^119^. The weighted average of the square of individual coefficients (loading score) for the top three principal components (weighted by the principal component’s eigenvalues) was used to calculate the contributions of individual variables. Compositional dissimilarities among restoration durations (beta diversity) were partitioned into replacement and richness difference components with the R package adespatial.

Microbe-metabolite correlations were calculated with the Spearman method. The correlations with the absolute Rho values > 0.3 and the p-values <10^-4^ were selected.

Correlation analysis results were visualized in R using the packages igraph (v1.4.1) and ggraph (v2.1.0).

### Species interaction network

To investigate the species interaction network, we first selected species presented in more than 28 of all soil samples (>10%; 1929 species) at the polyculture site.

1. Spearman’s correlation coefficients |r| > 0.6 and *p.adj* < 0.01 were filtered to construct networks, which is comparable to previous studies^120^. We then extracted each group’s sub-networks based on the species’ presence (abundance >0).
2. Various indexes, including density, negative percentage, average degree (connectivity), average path length, diameter, clustering coefficient (transitivity), eigenvector centrality, betweenness centrality, closeness centrality, degree centrality, and natural connectivity^121^ were calculated to describe the overall topologies of species interaction-networks in different regions. And degree, betweenness, local clustering coefficient, closeness, and page rank were calculated to indicate the properties of individual nodes in the network.
3. To estimate the level of modularity and the number of modules in different networks, we used a fast greedy modularity optimization algorithm^122^. Each module is defined as a group of species more closely connected than species outside of the module.
4. The within-module degree Zi and participation coefficient Pi were calculated to define the keystone taxa that may have special topological roles in the network^123^; Zi measures how ‘well-connected’ node i is to other nodes in the module while Pi measures how ‘well-distributed’ the links of node i are among different modules. They are calculated as follows: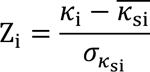
5. Where *K*_i_ is the number of links of node i to other nodes in its module si, *K*_si_ is the average of k over all the nodes in si, and σ_Ksi_ is the standard deviation of *κ* in si.
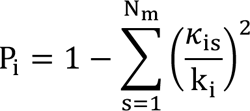 where *K*_is_ is the number of links of node i to nodes in module s, and k_i_ is the total degree of node i.
6. Invulnerability tests of networks calculate the natural connectivity to evaluate the stability of networks. Specifically, natural connectivity was calculated each time after removing five nodes from a network until 30% of nodes remained. The downtrend level of natural connectivity indicated the connectivity performance of the network after being damaged to a certain extent.

Cohesion was calculated to quantify the connectivity of microbial communities in each group. Cohesion contains both positive and negative cohesion values, which indicate that associations between taxa attributed to positive and negative species interactions and similarities and differences in niches of microbial taxa^124^. The pairwise Pearson correlation matrix across taxa was calculated using the abundance-weighted matrix. After bootstrapping taxa with 200 simulations, average positive and negative correlations were calculated to generate a connectedness matrix. Finally, positive and negative cohesions were calculated for each sample by multiplying the abundance-weighted and connectedness matrices. The absolute value of negative: positive cohesion is an important index for community stability. The formulas are as follows: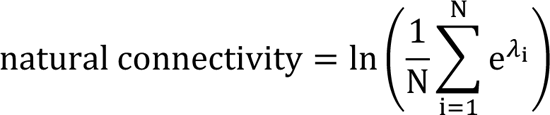

Where N denotes the number of nodes of the network, λ_%_ represents the eigenvalue of the network adjacency matrix.
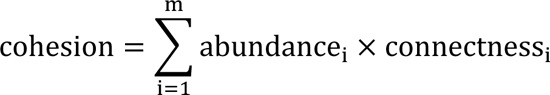
where m is the total number of taxa in a community.

All network analyses were carried out by the R programming mainly based on igraph packages (v1.3.5). Networks were visualized with the igraph package and the interactive platform Gephi.

### Construction of AEMs

AEM (asymmetric eigenvector maps) is an eigenfunction method originally developed to model multivariate (e.g., species) spatial distributions generated by an asymmetric, directional physical process. AEM is suitable for the temporal series analysis because the processes associated with time are directional: changes occur from time point 1 to time point 2, not the reverse^125^. AEM eigenfunctions are calculated from the temporal neighborhood matrix and temporal weighting matrix derived from the restoration duration data. Each AEM eigenfunction independently shows a broad- or fine-scale temporal profile in the ecological data and can be directly visualized as curve graphs. This study used the AEM, ade4, mvpart, vegan, magrittr, and packfor packages to construct the AEM eigenfunctions at the polyculture site. The AEM eigenfunctions were forward-selected on the taxonomy abundance (genus level) data using redundancy analysis (RDA) to remove insignificant AEM eigenfunctions. We kept the significant AEM variables for use in the following variation partitioning assessments.

### Variation Partitioning analysis

Overall variation partitioning was performed as previously described^23^. In brief, we used distance-based redundancy analysis (dbRDA) to conduct constrained ordinations on the genus dataset. We collected and prepared 46 or 44 metadata variables for each sample in the SI and DI regions, respectively. These variables were divided into “physicochemical”, “temporal”, and “technical” groups, and variables were forward-selected within each group. In both SI and DI regions, the correlation between each pair of groups was either negligible (low R^2^) or insignificant (*p* > 0.05). For instance, in the SI regions, the Mantel test with 999 permutations and “Bray-Curtis” distance showed R^2^ = 0.00637, *p* = 0.291 between “physicochemical” and “temporal” groups; R^2^ = 0.135, *p* = 0.001 between “physicochemical” and “technical” groups; R^2^ = 0.0227, *p* = 0.018 between “technical” and “temporal” groups. The varpart function of the vegan package was used to partition the variation (based on squared Bray-Curtis distance) of the dataset based on the three defined groups of variables. Adonis, an improved PERMANOVA analysis, was used to evaluate the variation explained by individual variables.

For genus-based variation partitioning in SI regions: we first performed a forward selection of all 46 variables (Table S1 and Figure S3F) over the abundance data of genera present in more than 100 samples (565 genera) to select the best representative variables. We then constructed multivariate linear regression models for 565 genera; 441 models had adjusted *p*-values less than 0.05. The formula is: asinh(CPM) ∼ batch + moisture + Mn + Ti + Ec + C:P + Ks + C:N + AEM1 + AEM23 + AEM20 + AEM2 + AEM47 + AEM4

For genus-based variation partitioning in DI regions: We first performed a forward selection of all 44 variables (Table S1 and Figure S9K) over the abundance data of genera present in more than 70 samples (579 genera) to select the best representative variables. We then constructed multivariate linear regression models for 579 genera; 215 models had adjusted *p*-values less than 0.05. The formula is: asinh(CPM) ∼ batch + moisture + Median size + AEM50 + AEM1 + AEM11 + AEM4

We then used the hierarchical partitioning method from the relaimpo package to evaluate the contributions of each variable via all potential ways of adding the terms in regression models (instead of evaluating the R^2^ based on one particular regression model). We chose not to evaluate R^2^ based on one particular regression model in this analysis because we wanted to assess the relative importance of each variable while taking into account all possible combinations of variables in the regression models. The contribution of each variable was calculated in conjunction with all other variables, and the contributions of each group (physicochemical, temporal, and technical) were evaluated by summing up the contributions of the variables in each group. It is important to note that each group’s total proportion of contributions is evaluated by the total explained variation of each model rather than the total variation. To visualize the group-based variation partitioning results, ternary plots were employed using the *ggtern* package.

To evaluate how soil properties impact the cohesion (Figure 3D), we used the *dredge* function in the R package Muti-Model Inference (MuMIn)^126^ to select the best-fitting models that minimized the AICc. Because the maximum number of predictors in the *dredge* function is 31, based on the previous regression model results, we selected 20 metadata grouped into biotic (Shannon diversity of bacterial and fungi, and FVC) and abiotic factors (C: N ratio, C:P ratio, Ti, Mn, EC, Ks, As, K, Ba, Cr, Ni, Mo, Cu, Mg, Zn, Fe, Al) variables. A multiple regression model was used to reveal the joint effects of these variables on cohesion. Before calculation, all predictors and response variables were standardized using the Z-score.

### Graph-based permutation test

To evaluate the impact of temporal variables on the soil microbiome, we calculated Bray-Curtis distances between samples at the genus level. We then constructed a nearest neighbor (NN) tree with the number of neighbors set as 1 (knn = 1), and colored each node in the tree by the duration of restoration. A “pure” connecting edge indicates that the two nodes (samples) connected are from the same group (restoration duration). In contrast, a “mixed” connecting edge indicates that the two nodes are from different groups. We counted the number of pure and mixed edges in the tree, then permuted all nodes’ labels while maintaining the initial tree structure. This was done to evaluate whether the observed number of pure edges is statistically significant over the background distribution^117^. The number of pure and mixed edges was counted in each permutation, and the background distribution of the number of pure and mixed edges for the initial tree was constructed (N = 9999). Finally, we evaluated the probability of observing the number of observed pure edges (or greater) in the initial tree based on the background distribution. It is important to note that we could not generate a p-value less than 1/N (or p = 0.0001) due to the permutation-based p-value estimation method. This means that the actual probability may be lower.

### Fuzzy c-means clustering

We used fuzzy c-means clustering (FCM)^127^, an unsupervised machine learning technique that divides the population into several groups or clusters, to explore the latent patterns of features across time or groups. They are similar if features are classified into the same group or cluster. Three methods (Elbow, Silhouette, and Gap statistic) were used to determine the optimal cluster number. The clustering results were further optimized through visualizations of the results (PCA and line plots). In FCM, the membership score is the probability of a feature belonging to a cluster, and we assign each feature to a cluster based on its top membership score. Only features with membership scores higher than specified values were considered and visualized in line plots. The alpha (transparency) of each line is directly related to the value of its membership score.

### The predictive modeling of the restoration duration

To predict the planting year based on the taxonomic profile across different domains of life, we referred to the R script^23^ pipeline with the glmnet package. The LASSO logistic regression classifier from the glmnet package was employed. It generates a classifier with a small number of selected features from the hundreds of genera in our dataset. These features are biologically interpretable as the linear combinations of variables in the logistic regression. The LASSO classifier also provides a realistic estimate of the generalized error during cross-validation, thanks to the built-in feature selection process. This contrasts the two-step method, where a supervised feature selection step is implemented before cross-validation, which can lead to over-optimistic accuracy estimates. Because only one of the top correlating features is selected before model training/testing, the LASSO classifier prevents information leakage from highly correlated features between the training and testing datasets.

Here is the complete workflow step by step:

1. Transforming taxonomic data with arcsinh, which behaves similarly to a log transform at high values, but is approximately linear near zero (unlike the log, it can handle zeros or small negative values);
2. To achieve the best balance between resolution and accuracy, genus-level datasets were used. The duration of 11, 12, 14, and 15 were selected from the SI dataset, except for the 17-year group, due to its high similarity to the 15-year group.
3. Partitioning our data randomly into 10 equal subsamples and then keeping one subsample for testing (one-fold) and using the remaining subsamples (nine-fold) for training through nested cross-validation with k = 10 folds. The train data was used to generate a hyperparameter for the selected model (in this case, the penalizing parameter lambda for feature selection). The test data was used to evaluate the hyperparameter for the selected model. The resulting model obtained from the train data was then evaluated on the test data and generated model performance parameters, such as the multi-class area under the curve (mAUC), accuracy, specificity, F1 score, etc. We resampled the entire dataset 10 times to generate 100 internal model performance parameters to assess its stability. Weights for samples in each planting year were adjusted for every training iteration to account for sample size variations in the four planting years;
4. Based on the average prediction scores of the resampled 10×10 internal testing, planting year prediction scores for each sample were obtained. Mean predicting scores of each planting year were used to calculate the multi-class ROC curve using one versus all approaches for each year;
5. For model interpretation, feature extraction, and visualization, non-zero coefficients for each year were examined and visualized as a heatmap. The square of the coefficient normalized by the total sum of the squared coefficients for each year was used to explain its contribution to the prediction model. Log-odds ratios of the respective features were represented as the bar length of the lollipop plot.

### Microbiome community assembly

To determine the potential importance of stochastic processes on community assembly based on neutral theory, we assessed the fit of the Sloan Neutral Community Model^128^ to species distributions. This neutral community model (NCM) predicts the relationship between species detection frequency and their relative abundance across the wider soil community. In this model, Nm is an estimate of dispersal between communities. The parameter Nm determines the correlation between occurrence frequency and regional relative abundance, with N describing the metacommunity size and m estimating the immigration rate^128^. The parameter R^2^ represents the overall fit to the neutral model.

To understand the assembly rules of microbial communities based on the niche-based theory and reveal factors that dominate the different assembly processes during restoration, we conducted the following analyses based on Feng et al.^129^ with minor modifications:

1. We focused on the bacteria annotated in all samples from SD and SI groups to build the phylogenetic tree based on the Genome Taxonomy Database, as bacteria occupied a major part of the taxonomic profile (1262/1416, 1416 species that existed in more than 10% samples).
2. Mantel correlograms with 999 randomizations using the R package microeco (0.11.0) were used to evaluate the phylogenetic signal across a range of phylogenetic distances. Significant positive correlations between bacterial niche differences and phylogenetic distances were strongest at short phylogenetic distances (approximately 0%-42% of the maximum phylogenetic distances), indicating that bacterial environmental preferences were phylogenetically conserved across relatively short phylogenetic distances. We then evaluated the between-community mean nearest taxon distance (βMNTD)^130^ and β-nearest taxon index (βNTI)^131^ to quantify phylogenetic turnover between communities, using the R packages NST (3.1.10), picante (1.8.2), and ape (5.6-1). These two indexes emphasize relatively short phylogenetic distances^131^, which is appropriate as the strong phylogenetic signal was observed across short phylogenetic distances.
3. Four ecological processes govern community assembly: homogeneous selection, variable selection, homogenizing dispersal, dispersal limitation, and a condition for which no single process dominates community assembly (referred to as “undominated”)^132^; We combined the outcome of βNTI with a second null model Bray-Curtis-based Raup-Crick (RCbray)^133^ to infer the relative influences of homogenizing dispersal and dispersal limitation. The detailed principle and calculation can be found at https://github.com/stegen/Stegen_etal_ISME_2013.

### Antibiotic resistance genes and virulent factors analysis

To annotate all quality-filtered unassembled sequencing reads to the antibiotic-resistance genes (ARGs), we used the Comprehensive Antibiotic Resistance Database (CARD, downloaded on August 10, 2021)^134^ by using the CARD RGI software (v.5.2.0) with the option “--aligner bwa”. The abundance of ARGs was calculated as mentioned for CAZymes.

We used the Virulence Factors Database (VFDB) VFDB_setB_pro.fas (27504 sequences, downloaded on June 6, 2022)^135^, which contains experimentally verified virulence factors, to annotate the VFs. We performed the alignment using DIAMOND with BLASTP (coverage > 50% and e-value < 1 x 10^-5^). The taxonomic source information for each protein was obtained from the overlap between VFDB_setB_pro.fas and our Kraken2 results, and we evaluated the abundance of VF using aCPM.

### CRISPR Array and Protospacers analysis

We extracted CRISPR arrays and spacers from assembled contigs from the polyculture site using metaCRT^136^ with default parameters, which yielded 54,251 CRISPR arrays and 204,909 spacers sequences associated with 53,694 contigs. We created spacer clusters with 80%, 90%, 95%, and 98% similarity thresholds using CD-HIT^137^ (4.8.1) with default settings resulting in sets of 65,692, 138,341, and 168,918, 182,373 cluster representatives, respectively. We assigned the taxonomic information to the host contigs of CRISPR arrays using kraken2^33^. To associate detected spacers within defined Viruses, a BLASTN (2.10.1+) search with the command line parameters-task blastn-short - qcov_hsp_perc 90-perc_identity 90 -max_hsps 3 -num_threads 12 -outfmt 6 was performed for the spacer set against the IMG/VR database^138^ (IMG_VR_2020-10-12_5.1 - IMG/VR v3). The hits with at least 90% sequence identity to a spacer and at least 90% sequence coverage were candidate protospacers. As a result, 79,920 spacer matches were detected in viral sequences. We filtered the mapping and retained matches with the top 10% bitscore. Since one spacer would match several protospacers, We parsed the LCA taxonomy for protospacers using TaxonKit^139^.

### Functional annotations of genes

To maximize functional predictions of genes, all quality-filtered unassembled sequencing reads were aligned to the NR database (downloaded on Jun 30, 2020) via BLASTP of DIAMOND (v2.0.14.152)^140^ with coverage > 50% and e-value < 1 × 10^-10^. We extracted the best hits for each metagenomic sequencing read and evaluated the abundance of proteins in each sample using aCPM.

We then aligned the NR database against the KEGG database (downloaded on November 8, 2021, with 35729277 genes) using BLASTP implemented in DIAMOND (coverage > 50% and e-value < 1 x 10^-5^) to generate the NR-KO index. Each gene was assigned to the KO by the highest-scoring annotated hit(s) containing at least one HSP scoring over 60 bits. We then searched the gene patterns from each sample against the NR-KO index and identified differentially enriched KO pathways according to their reporter scores^141^. The taxonomic source for genes or metabolic markers was based on the overlap between the NR database and our Krakrn2 results.

For focused functional annotations, the NR database was aligned against the NCycDB^66^, MCycDB^60^, SCycDB^142^, and PCycDB^143^ databases to generate a functional index using the BLASTP implemented in DIAMOND, with parameters set to coverage > 50% and an e-value cutoff of 1 × 10^-5^. We then searched the gene profiles from each sample against the generated functional index. Integrated with the id mapping file and the abundance of each protein, we obtained the functional profiles of nitrogen cycle, methane cycle, sulfur cycle, and phosphorus cycle. Based on the taxa information from NR and Kraken2 annotation results, we determined the source taxon for each gene in every sample, resulting in gene-function-taxa-abundance profiles.

To identify carbohydrate-active enzymes (CAZymes)^59^, the NR database was first compared against the dbCAN (released on July 31, 2019) database^144^ using DIAMOND with BLASTP (coverage > 50% and e-value < 1 x 10^-5^) to generate a CAZymes index. The protein profiles of each sample were further annotated with the CAZymes index. Subsequently, for any sample N, the abundance of CAZymes was calculated as follows:

Step 1: Calculate the copy number of each gene:
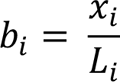

Step 2: Calculate the relative abundance of gene i:
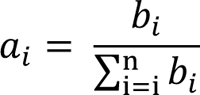

*x*_i_: the number of mapped genes in sample N

*L*_i_: the length of gene i

*b*_i_: the copy number of gene i from sample N

*a*_i_: the relative abundance of gene i in sample N

Enzymes involved in degrading cellulose, hemicellulose, xylan, and lichenin were derived from Glycoside Hydrolases (GHs, hydrolysis, and/or rearrangement of glycosidic bonds), Polysaccharide Lyases (PLs, non-hydrolytic cleavage of glycosidic bonds), Carbohydrate Esterases (CEs, hydrolysis of carbohydrate esters), and Auxiliary Activities (AAs, redox enzymes that act in conjunction with CAZymes) using customized python scripts based on the dbCAN database.

The integrated taxon-function maptrees were visualized in R using devtools (2.4.3), igraph (1.3.1), tidyverse (1.3.1), viridis (0.6.2), data.tree (1.0.0), phyloseq (1.38.0)^48^, ggtree (3.2.1), and ggraph (2.0.5) packages.

### Reporter score functional enrichment analysis

The Reporter score functional enrichment analysis is performed using the “directed” mode of the ReporterScore package (https://github.com/Asa12138/ReporterScore). This method is based on the assumption shared by many differential analysis methods, which states that the abundance of most KOs does not significantly change. The specific steps involved in this method are as follows:

1. Wilcoxon Rank Sum Test: Use the Wilcoxon rank sum test to determine the significance (*P* value) of the difference in abundance for each KO between two groups (represented as P_koi_, where i refers to a specific KO). Divide each P value by 2 ((*P*_koi_ = *P*_koi_/2), transforming the range from (0,1] to (0,0.5].
2. Conversion to Z values: Convert the *P* value of each KO into a Z value (Z_koi_) using the inverse normal distribution formula:
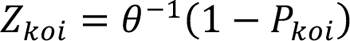
 Since the P value is less than 0.5, all Z values will be greater than 0.
3. Determination of Up-regulation or Down-regulation: Consider whether each KO is up-regulated or down-regulated. Calculate Δ*KO*_i_:
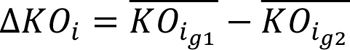
 *KO*_ig1_ represents the average abundance of *KO*_ig2_ in group1, *KO*_i_ represents the average abundance of *KO*_ig2_ in group2. Then:
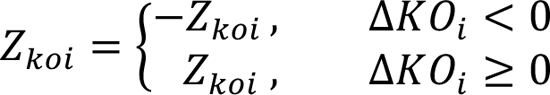
 *Z*_koi_ greater than 0 indicates up-regulation, while *Z*_koi_ less than 0 indicates down-regulation.
4. Pathway-level Z value calculation: Aggregate the Z_koi values to obtain the Z value for each pathway. The formula is as follows:
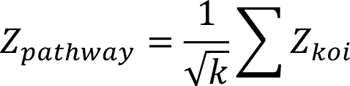
where k represents the total number of KOs annotated to the corresponding pathway.
5. Significance Evaluation: Perform permutation (1000 times) to obtain a random distribution of Z*_pathway_*. Calculate the adjusted Z value for the pathway using the formula:
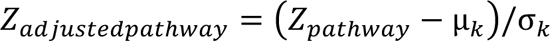
where μ_k_ is the mean of the random distribution and σ_k_ is its standard deviation. The resulting *Z_adjustedpathway_* is the Reporter score value enriched for each pathway. In this “directed” mode, the Reporter score indicates the direction of enrichment. A larger positive value indicates significant up-regulation enrichment, while a smaller negative value indicates significant down-regulation enrichment.

### Gradient boosting regression model

We implemented the gradient-boosting regression model to predict key soil health properties using microorganisms and metabolites while selecting the top important variables. Here is the step-by-step workflow:

1. We focused on the 878 genera and 1499 metabolites annotated in over 10% of SD, DI, and SI samples to construct the Gradient boosting regression model.
2. Ks and carbon levels were selected as crucial indicators for ecosystem restoration.
3. We partitioned our data randomly into 10 equal subsamples, reserving two subsamples for testing (two-fold) and employing the remaining subsamples (eight-fold) for training. The training data generated a gradient-boosting model through cross-validation with 10 folds. The test data was employed to evaluate the model’s performance. The resulting model obtained from the train data was then assessed on the test data, generating performance parameters such as R and *p-values* by permutation-based estimation method (N = 999). We resampled the entire dataset 10 times to evaluate its stability. The top 30 relative influence variables from the 10 models were chosen to construct the simplified gradient-boosting regression model as described. GLM was used to evaluate the correlations between the selected variables and Ks or carbon.

### Structural equation modeling

Structural equation modeling (SEM) was applied to determine the direct and indirect contributions of restoration duration and fungal diversity to soil nutrients. SEM analysis was conducted with R package lavaan (v0.6-12). The model’s fitness was examined by a chi-squared test, comparative fit index (CFI), and the root mean square error of approximation (RMSEA).

### Bi-directional mediation analysis

Bi-directional mediation analyses were performed using the *mediate* function from the mediation R package (v4.5.0) to investigate the causal role of the microbiome in shaping soil properties through annotated metabolites (n = 749). The bootstrap number was set as 199. The *p-values* were adjusted by the FDR method. We then selected linkages with *p.adj* < 0.05. Next, we checked whether the soil properties were associated with metabolites and microbes whether the microbes were associated with metabolites by Spearman tests. Finally, we checked the *p-values* of reverse mediation analysis and selected mediation linkages with *p-values* > 0.05.

### Population genetics analysis

For population genetics analysis, we generated a list of the top 100 abundant reference genomes based on Kraken2 species profiles. A collection of these genomes was used as the reference to map the reads sample by sample using ngless (v1.5.0)^145^. Within ngless, mapping was done by BWA-MEM and followed with alignment filtering, which only kept alignments with at least 97% identity and 45 bp length and discarded multi-mapped reads. Then, we merged the resulting bam files using the ‘merge’ command of samtools (v1.9). We used metaSNV ^146^ to estimate SNP density (the value of SNPs/kbps) and nucleotide diversity (π) based on the output SNP profiles.

To calculate the evolutionary metrics for each sample, we used inStrain^147^. Specifically, we selected the top 30 abundant species in Kraken2 profiles as the reference genome set for inStrain profiling. Next, we used bowtie2 (v2.4.2) to map reads to the reference genome set sample by sample. Then, using the ‘profile’ command of inStrain, we generated the microdiveristy profiles for each sample. To keep species with sufficient coverage in individual samples, we filtered out the species with < 0.1 breadth_minCov (meaning that at least 10% of the bases in the genome were covered by at least five reads, as minCov was set as 5). We called genes of the top 30 species using prodigal (v2.6.3), whose output can be used by inStrain to profile the microdiveristy at the gene level. The genes with > 0.5 breadth_minCov were considered present. The estimation of gene pN/pS was implemented by inStrain. Genes with pN/pS < 1 were undergone positive selection. Genome-wide pN/pS were calculated by taking SNVs of all genes across a genome. Genes were also annotated by Diamond (v2.0.15) with the manually constructed reference database of biogeochemical cycling genes. The pN/pS of biogeochemical cycles were the average pN/pS of the involved genes.

