## Supplemental information for "Progressive community, biogeochemical and evolutionary remodeling of the soil microbiome underpins long-term desert ecosystem restoration"

### Supplementary Text

#### The phosphorus and sulfur cycles of the desert soil microbiome during ecosystems restoration

Sulfur is a vital macronutrient required for growth and functioning. Organic sulfur accounts for approximately 95% of total sulfur in soils in two major forms: sulfate-esters and sulfonates. These organic sulfur forms are not directly available to plants, critically dependent upon microbial mineralization to sulfate ($SO_4^{2-}$)[1]. We annotated 200 genes in the sulfur cycle by the SCycDB[2]. The sulfur oxidation (*tsd AB and doxD*), reduction (*shyC*), and dissimilatory $SO_4^{2-}$reduction (*qmoB*) increased over time (Figure S7G). Microbial sulfur oxidation and reduction are the most active and important metabolic processes in the sulfur cycle. The efficiency of sulfur oxidation increases at higher soil temperatures, in moist soils, soils with higher pH, and higher amounts of soil organic matter and availability of other nutrients[3,4]. Microbial $SO_4^{2-}$ reduction includes assimilatory and dissimilatory $SO_4^{2-}$ reduction. The assimilatory reduction pathway incorporates reduced sulfur into sulfur-containing cell components such as cysteine and methionine. In contrast, in dissimilatory reduction, $SO_4^{2-}$ is reduced to inorganic sulfide to obtain energy. The relative abundance of the dissimilatory reduction pathway increased, while no differences in assimilatory reduction abundance were observed (Figure S7G). Previous research has suggested that sulfur uptake and assimilation are derepressed under sulfur starvation or high demand for sulfur metabolites[4]. The stable assimilatory reduction abundance suggests that the soils are not sulfur-deprived or that the assimilatory sulfate reduction would decrease. The inorganic and organic sulfur transformation and organic sulfur transformation increased (Figure S7G), indicating greater sulfur immobilization and mineralization activation. The genes involved in sulfur oxidation, sulfur reduction, and dissimilatory sulfate reduction all had a bacterial source (Figures S7H-S7J). All four kingdoms contributed to the dynamic inorganic and organic sulfur transformation, with a major source from Proteobacteria and Actinobacteria (Figure S7K). Actinobacteria were the key drivers of the dynamic organic sulfur transformation (Figure S7L), and all related phyla participated in organic sulfur transformation (Figure S7M).

Notably, the SD control also experienced an overall decrease in the sulfur cycle (Figure S6B), and no dynamic change in the sulfur cycle in the DI regions was observed (Figure S10H).

Phosphorus is essential for all living organisms' energy metabolism, genetic materials, carbon metabolism, metabolic signaling, and cell structures[5]. The phosphorus cycle is unique and different from the carbon and nitrogen cycles because phosphorus does not exist in a gaseous form, making it the limiting nutrient in natural terrestrial ecosystems[6]. It has long been recognized as a critical determinant of crop productivity and the growth of microbes[7]. Soil phosphorus comprises organic and inorganic forms, with 30-65% of phosphorus in organic forms unavailable to plants. Soil microorganisms play a crucial role in altering soil phosphorus availability by processing and transforming organic forms of phosphorus (such as plant residues, manures, and microbial tissues) into plant-available forms through mineralization and solubilization. We annotated 140 genes in phosphorus cycling by the PCycDB[8]. The differential phosphorus cycling genes over time are involved in extracellular phosphorus transport and intracellular phosphorus metabolism (Figure S8A). The two-component system, transporters, and organic phosphoesters hydrolysis are major microbial processes to regulate, transport, and uptake phosphorus sources from the environment. The two-component system decreases over time ($p$ = 0.08; e.g., *phoP*, *senX3*, and *regX3*), indicating a deficiency in the extracellular inorganic phosphorus (Figure S8A).

Phosphorus cycling is closely linked to carbon and nitrogen dynamics in the soil. Phosphorus is essential for photosynthetic carbon metabolism and the production of photo-assimilates. As a structural component of various sugar phosphates such as ribulose-1,5-bisphosphate (RuBP) and fructose 6-phosphate, phosphorus limitation impairs the regeneration of RuBP, which reduces rubisco activity and $CO_2$ fixation[9,10]. Additionally, the regeneration of RuBP in chloroplasts is an ATP-dependent process, and phosphorus-deficient plants have a very low pool of ATP[11]. Many studies have reported the critical nutrient role of phosphorus in biological $N_2$ fixation in leguminous plants[12,13]. Phosphorus deficiency significantly impairs nodulation and nodule functioning in soybean plants[12,13]. The $N_2$ fixation process is highly energy-demanding and requires at least 16 moles of ATP to reduce one mole of $N_2$. In this study, the increase in the more efficient $CO_2$ fixation pathway of the HP/HB cycle (Figure 5A) and the non-significant change in $N_2$ fixation (Figure 5E) may be linked to the scarcity of phosphorus in the soil. Although we observed an increase in total phosphorus concentration over time (Figure 1D), there is still a significant gap compared to the most common natural, unfertilized, and uncultivated topsoil (20 to 1000 mg/kg)[14]. Therefore, abundant phosphorus accumulation is necessary for more activated carbon and nitrogen cycling during restoration.

Intracellular inorganic phosphorus is required for phosphorylation reactions within the central carbon metabolism and the eventual generation of precursors for the biosynthesis of organic phosphorus biopolymers: nucleic acids and phospholipids (PCYdb). Here we follow the incorporation of inorganic phosphorus (Figure S8A), starting with the initial catabolism of sugar, a common carbohydrate in soils, and the derivative of cellulose and hemicellulose. Sugar is phosphorylated to produce sugar-phosphate via the phosphoenolpyruvate-dependent Phosphotransferase system (Figure S8A). The increased phosphotransferase system (*ptsH*) provided more sugar-phosphate, subsequently increasing the pentose phosphate pathway (Figure S8A; *prsA* and *phnN*). Meanwhile, the increased phosphonate and phosphinate metabolism (*phnO*) generated more intermediates for the pentose phosphate pathway (Figure S8A). Within the pentose phosphate pathway, sugar-phosphate produces ribose-5-phosphate (R5P) through oxidation and decarboxylation reactions (Figure S8A). α-D-ribose-1-diphosphate-5P (PRPP), a key phosphonate precursor in the biosynthesis of the nucleotides (i.e., purine and pyrimidine), the amino acids (i.e., histidine and tryptophan), and the cofactors (i.e., NAD and NADP), could be synthesized by ribose-phosphate pyrophosphokinase encoded by prsA through R5P and ribose 1,5-bisphosphokinase *(phnN*) via ribose-1-5-phosphate (Figure S8A). Both purine and pyrimidine nucleotides are essential to biomass growth as they are building blocks for RNA, DNA, and the energy carrier molecules (ATP,

NAD, and NADP; Figure S8A). The increasing pentose phosphate pathway and pyrimidine metabolism (*cmK* and *nrdEF*) suggested more activated microorganisms' synthetic ability over time (Figure S8A). The genes in the phosphotransferase system were mainly encoded by Actinobacteria and Proteobacteria (Figure S8B). Proteobacteria was the major phyla responsible for enhancing phosphonate and phosphinate metabolism, and genes encoded by archaea all increased (Figure S8C). However, the genes that participate in pyrimidine metabolism with a source from archaea all decreased over time (Figure S8D). The genes involved in pyrimidine metabolism were mainly from Actinobacteria, Bacteroidetes, Proteobacteria, and Planctomycetes (Figure S8D). All the related phyla participate in pyrimidine metabolism, as RNA and DNA are the genetic material in all living organisms (Figure S8E).

In the SD control, the lowest phosphotransferase system may be responsible for the lowest phosphonate and phosphinate metabolism and pyrimidine metabolism, as decreased substrate sources were provided for the following biological reactions (Figures S6A and S6B), suggesting the overall inactive phosphorus turnover in SD. There was no dynamic change in the phosphorus cycle in the DI regions (Figure S10H). In short, the phosphorus and sulfur cycles were highly activated in the SI region during restoration.

### Supplementary Figure Legends

**Figure S1. Representative photos and overview of soil health properties of the polyculture and monoculture sites, related to Figure 1.**
(A) Map showing the general geographic regions around the two restoration sites.
(B) Schematic drawing and aerial views of the polyculture site. Light green indicates natural recovery with aerial seeding, and dark green represents planting regions with drip

irrigation (DI) or manual subsurface irrigation (SI). The restoration site was divided into two regions (orange and purple). The lines ended with arrows and dots indicate the sampling lanes in each region.
(C) Representative pictures of each sampling region of the polyculture site.
(D) The sampling sites and representative pictures of the monoculture site. Samples were collected from the biocrust and non-biocrust regions (Picture on the right), as well as the natural recovery region (NR) and sand dunes (SD2).
(E) Fractional vegetation cover (FVC) index of the monoculture site. The inset boxplot represents the FVC index of different groups. Different lowercase letters indicate $p < 0.05$.
(F) Heatmap of soil physical, chemical, and biological properties. The properties are colored yellow (increasing), green (decreasing), or black (non-significant) based on significance ($p.\text{adj} < 0.05$) by the GLM in the SI regions. SD, sand dune; Numbers from 4-17 indicate restoration duration. Ks, hydraulic conductivity; FVC, fractional vegetation cover.

**Figure S2. Overview of the study at the polyculture and monoculture sites, related to Figure 1.**
(A) Representative differentially abundant soil health attributes on the polyculture site (Kruskal-Wallis; $p.adj < 0.05$). Different lowercase letters indicate $p < 0.05$. Notably, all samples belong to the "Non-saline" category ($0 < EC < 2000$ μS/cm).
(B) Effects of the duration of restoration (4 – 9 years) on soil health properties in DI regions (GLM; $p.adj < 0.05$). Ks, hydraulic conductivity; FVC, fractional vegetation cover.
(C) Effect of the duration of restoration (0.5 – 6 years) on soil health properties in non-biocrust (NBC) and biocrust (BC) regions (GLM; $p.adj < 0.05$). Ks, hydraulic conductivity.
(D) Differentially abundant soil health attributes in the monoculture site. The numbers of samples in each group are SD2 (7), NBC (52), BC (20), and NR (5). Different lower-case letters indicate $p < 0.05$ by the Kruskal-Wallis.
(E) Density plot of read counts per sample for DNA in the polyculture and monoculture sites.
(F) Density plot of total species abundance based on aCPM per sample for DNA in the polyculture and monoculture sites.
(G) Randomized species accumulation curves for the polyculture (left panel) and monoculture (right panel) sites.
(H) Estimated completeness and contamination of 1565 metagenome-assembled genomes (MAGs) recovered from 382 desert metagenomes at two sites. High-quality MAGs (completeness > 90%; contamination < 5%) are shown in red, medium-quality genomes (completeness ≥50%; contamination < 10%) in green, and Low-quality genomes (contamination < 10%) in blue. Density plots along the x and y axes show the distribution of completeness and contamination, respectively.
(I) Phylogenetic trees of archaeal metagenome-assembled genomes (MAGs) of two sites. Branches and inner rings are colored by kingdoms. The intermediate ring labels the quality of MAGs, with red, green, and blue representing high-, medium-, and low-quality MAGs. The outer ring denotes the relative abundance of MAGs.
(J) The number of metagenome-assembled genomes (MAGs; top), species-level genome bins (SGBs; middle), and the unknown SGB (uSGB; bottom) percentage in each phylum

of two sites. Two phyla are from Archaea, and the others belong to Bacteria. The SGBs without an existing reference genome (could not be annotated at the species level by GTDB-tk) were defined as unknown SGBs (uSGBs), while the SGBs having at least one MAG could be annotated at the species level as known SGBs (kSGBs).

**Figure S3. The source of variation of the highly dynamic desert soil microbiome in the SI regions, related to Figure 2.**
(A) Shannon indexes of different domains of life in different regions of the SD control and SI regions. Different lowercase letters indicate $p < 0.05$ by the Kruskal-Wallis. The significant *p*-values of SI samples based on GLM are labeled.
(B) Detrended Correspondence Analysis (DCA) of all microbiota data. The lengths for DCA1 and DCA2 axes are 1.594 and 1.308, respectively. This indicates the soil microbiome data's environmental gradients are short and linear. Henceforth, PCoA, PCA, RDA, or dbRDA methods can be used.
(C) PCoA analysis of samples with Bray-Curtis distance showing that microbiome composition transited along the duration of restoration ($P < 0.001$, adonis). The colors of nodes represent samples of different restoration duration.
(D) Ternary plots of beta diversity comparisons (using the Sørensen dissimilarity index) for different kingdoms. Each point represents a pair of sites and 7021 pairs of sites are included. The position of each dot is determined by a combination of values of the S (similarity), Repl (replacement), and RichDiff (richness difference) matrices. The mean values of S, Repl, and RichDiff are shown below each plot.
(E) Plotting of asymmetric eigenvector maps (AEM) variables over time. AEM1 had the lowest *p*-values and increased over time, suggesting that AEM1 represents the principal temporal trend of microbiota during restoration in SI regions.
(F) The Adonis analysis of forward-selected variables used in the dbRDA analysis. Different colors indicate membership of corresponding groups: cyan, temporal factors; yellow, physicochemical factors; gray, batch factor. AEM variables are listed in the order of (E).
(G) Variation partition using forward-selected variables (top) or all variables (bottom); these numbers were used to plot Figure 2F. Theoretically, including all variables in dbRDA analysis would provide the most optimistic estimation of total explained variation at the cost of statistical power to assess individual group/variable contributions due to multicollinearity.
(H) Heatmap of features selected by the regularized multi-class logistic regression model. Lollipop charts and the percentage information indicate the relative importance of each genus. The duration of 11, 12, 14, and 15 years were selected from the SI dataset.
(I) Stable internal performance of the restoration duration-predictive model (resampling ten times).
(J) Macro-average performance metrics from the resampling data (10 times).
(K) Abundance profiles of representative genera selected for restoration duration prediction. The *p.adj* are directly displayed.

**Figure S4. The assembly process, dynamic stability of the desert soil microbiome community, and functional diversity of metabolites at the polyculture site, related to Figure 3.**
(A) Neutral models for bacteria in the SD control and SI regions. The predicted occurrence frequency is shown as a solid black line, and dashed black lines represent 95% confidence intervals; green and red dots indicate the species that occurred more and less frequently than the prediction; Nm indicates the metacommunity size times immigration; m indicates the immigration rate; $R^2$ indicates the fit to the neutral model.
(B) Phylogenetic mantel correlogram showing significant correlational signals across short phylogenetic distances. Solid and open symbols denote significant ($p < 0.05$) and insignificant ($p > 0.05$) correlations, respectively, relating between-species niche differences to between-species phylogenetic distances.
(C) Key characteristics of the species interaction networks. The dynamic characteristics with *p.adj* < 0.05 by the GLM are labeled directly.
(D) The heatmap of module hubs and connectors. The presence (red) and absence (gray) of keystone taxa are shown for each group of samples.
(E) Network of all observed $MS^2$ features at the polyculture site, with node colors indicating superclass and edge colors indicating the connection types between the features.
(F) The circle plot illustrates the type and number of connections for all experimentally observed $MS^2$ features at the polyculture site. The inner bar plot represents the dynamically changed metabolite proportion based on fuzzy c-means with membership > 0.65 in the SI regions. The inner ring indicates the number of metabolites belonging to a certain type of connection, and the outer ring shows the connection types.
(G) The relative abundance of the top 8 superclass-level metabolites at the polyculture site.
(H) Venn plot showing the dynamically changed metabolites associated with duration of restoration identified by fuzzy c-means and GAM in the SI regions. Membership ≥ 0.75 by c-means or $p < 0.03$ by GAM.
(I) The top significantly changed metabolites associated with restoration duration in the SI regions. The GLM statistics are shown. Related parent 1, parent 2, or subclass information were labeled. HCAD, Hydroxycinnamic acids and derivatives.
(J) The functional pathways of the significantly changed metabolites in the NetID network analysis (abundance indicated by circle size) of SI regions. Functional categories (outer circles) and involved pathways (inner circles) are labeled in the SI regions. Metabolites are represented by filled circles and are colored red (increasing) or green (decreasing).
(K) Line plots of asymmetric eigenvector maps (AEM) variables of the SI regions over time.

**Figure S5. Functional diversity of desert soil microbiome in the SI regions, related to Figure 4.**
(A) Absolute abundance of differentially abundant KO pathways based on GLM over time.
(B) Reporter score bar plot comparing KO pathways enriched in the 11-year group compared with the 17-year group. Pathways having a reporter score greater than 1.6 in at least one group are plotted.

(C-D) Pie plot showing the proportion of different antibiotic-resistant mechanisms (C) and the abundance of top 10 ARG classes (D) in the polyculture site.
(E) PCA analysis of ARGs in the polyculture site.
(F) The total relative abundance of different VFs categories in the polyculture site.
(G) PCA analysis of VFs in the polyculture site. Colors indicate grouping information, and the shapes of points indicate different regions.
(H) Representative differentially abundant VFs categories. The GLM statistics are shown. PTM, post-translation modification.
(I-K) Sankey diagram showing the linkages between VFs, VFs categories, the corresponding species at the family level, and the corresponding species at the phylum level. (I), SD group; (J) the 17-year group with a focus on 'Adherence'; (K) the 17-year group with a focus on 'post-translation modification' and 'Stress survival".
(L) The significantly changed VF categories in the SI regions over time (GLM; $p.adj < 0.05$).
(M) The heatmap of changing VFs in SI regions (GLM; $p.adj < 0.05$). Among them, 15/26 and 9/16 VFs in increasing and decreasing patterns can be assigned taxonomic sources based on the VFs database and annotations from Kraken2.
(N) Redundancy analysis (RDA) of the relationship between the soil health variables and the relative abundance of VFs in the SI regions, a significant temporal trend could be observed.

**Figure S6. Nutrient cycling during restoration at the polyculture site, related to Figure 5.**
(A) Heatmap of the relative abundance of carbon (C), nitrogen (N), phosphorus (P), sulfur (S), and methane ($CH_4$) cycles.
(B) The boxplots of differentially active pathways (Kruskal-Wallis; $p.adj < 0.5$) in the polyculture site. The different letter above each boxplot denotes $p < 0.05$. HP/HB cycle, Hydroxypropionate-hydroxybutyrate cycle; DC/HB cycle, Dicarboxylate-hydroxybutyrate cycle; PPM, Phosphonate and phosphinate metabolism; IOST, Inorganic and organic sulfur transformation.
(C) Phylogenetic tree of species involved in the carbon cycle. Branch colors denote significantly temporally increasing (yellow) and decreasing (blue) species (GLM; $p.adj < 0.05$). The inner ring indicates the phylum and the remaining rings indicate the presence of the three carbon cycling pathways.
(D) Relative abundance of carbohydrate-active enzymes (CAZYmes) in the SI regions. The GLM statistics are shown. GHs, glycoside hydrolases; GTs, glycosyltransferases; CEs, carbohydrate esterases; PLs, polysaccharide lyases; AA, auxiliary activities; CBMs, carbohydrate-binding modules.
(E) The integrated taxon-function maptree of microbiome members encoding cellulase, hemicellulase, xylanase, and lichenase in SI regions. Outmost to innermost circles indicate the cellular organisms, kingdom, phylum, and class levels. The filled circles represent enzymes that are temporally increasing (red) or decreasing (green) (GLM; $p.adj < 0.05$). Circle size indicates the taxon-level average relative CPM abundance in the SI region. The phyla with the top relative abundance are labeled.
(F) Phylogenetic tree of species that encode carbohydrate-active enzymes in the SI regions. Branches are colored by increasing (orange) and decreasing (blue) trends (GLM;

*p.adj* < 0.05). The inner ring indicates the phylum information, and the rest rings indicate the presence of carbohydrate-active enzymes and are colored as in (E).

**Figure S7. The dynamic methane and sulfur cycles in the SI regions, related to Figure 5.**

(A) The simplified diagram of the methane cycle in the SI regions. The increasing (orange) and decreasing (blue) enzymes during restoration are listed (GLM; *p.adj* < 0.05). P-values are directly displayed for each pathway with increasing (orange) and decreasing (blue) trends. The box color of the enzymes represents which pathways the enzymes are involved in.

(B-E) The integrated taxon-function maptree of microbiome members involved in the anaerobic oxidation of methane (B), aerobic oxidation of methane (C), hydrogenotrophic methanogenesis (D), and methylotrophic methanogenesis (E). Outmost to innermost circles indicate the cellular organisms, kingdom, phylum, and class levels. The filled circles represent enzymes that are temporally increasing (red) or decreasing (green) as labeled in (A). Circle size indicates the taxon-level average relative CPM abundance in the SI region. The phyla with the top relative abundance are labeled.

(F) Phylogenetic tree of species that encode enzymes of the methane cycle. Branches are colored by increasing (orange) and decreasing (blue) trends. The inner ring indicates the phylum information, and the rest rings denote the presence of four pathways in (B-E).

(G) The simplified diagram of the sulfur cycle. The increasing (orange) and decreasing (blue) enzymes during restoration are listed (GLM; *p.adj* < 0.05). P-values are directly displayed for each pathway with increasing (orange) and decreasing (blue) trends.

(H-L) The integrated taxon-function maptree of microbiome members participating in sulfur oxidation (H), sulfur reduction (I), dissimilatory sulfate reduction (J), inorganic and organic sulfur transformation (K), and organic sulfur transformation (L). The colors of the circles and nodes are labeled as in (B).

(M) Phylogenetic tree of species that participate in the sulfur cycle. Branches are colored by increasing (orange) and decreasing (blue) trends. The inner ring indicates the phylum information, and the rest rings indicate the presence of five pathways and are colored as in (G).

**Figure S8. The dynamic phosphorus and nitrogen in the SI regions, related to Figure 5.**

(A) The simplified diagram of the phosphorus cycle. The increasing (orange) and decreasing (blue) enzymes during restoration are listed (GLM; *p.adj* < 0.05). P-values are directly displayed for each pathway with increasing (orange) and decreasing (blue) trends. The box colors of the enzymes represent the pathways.

(B, C, and D) The integrated taxon-function maptree of the microbiome members that involved in the phosphotransferase system (B), phosphonate and phosphinate metabolism (C), and pyrimidine metabolism (D). Outmost to innermost circles indicate the cellular organisms, kingdom, phylum, and class levels. The filled circles represent enzymes that are temporally increasing (red) or decreasing (green), as labeled in (A). Circle size indicates the taxon-level average relative CPM abundance. The phyla (black text; light blue circles) with the top relative abundance are labeled.

(E) Phylogenetic tree of species that encode enzymes of the phosphorus cycle. Branches are colored by increasing (orange) and decreasing (blue) trends. The inner ring indicates the phylum information, and the rest rings indicate the presence of three pathways and are colored as in (A).
(F-J) The integrated taxon-function maptree of microbiome members encode enzymes of the nitrogen cycle involved in the assimilatory nitrate reduction (F), denitrification (G), nitrification (H), anammox (I), and organic nitrogen metabolism (J). The detailed information is the same as in (B). The classes (purple text; light purple circles) with the top relative abundance are labeled in (F).
(K) Phylogenetic tree of species that encode enzymes of the nitrogen cycle. Branches are colored by increasing (orange) and decreasing (blue) trends. The inner ring indicates the phylum information; the rest rings denote five pathways in (F-J).

**Figure S9. The highly dynamic and the source of variation of desert soil microbiome at the DI regions, related to Figure 2.**
(A) Shannon indexes of different domains of life in different regions of the SD control and DI regions. Different lowercase letters indicate $p < 0.05$. The significant *p*-values of DI samples based on GLM are labeled.
(B) Significantly changed phyla (GLM; $p.adj < 0.1$).
(C) Fuzzy c-means clustering of the genera abundance profiles. The four potential clusters are shown.
(D) PCA analysis of genera abundance profiles, color-coded by clustering information from (C).
(E) The phylogenetic tree of significantly changed species over time. Only species with temporal patterns, estimated by GLM ($p.adj < 0.05$), spearman correlation ($p.adj < 0.05$), or C-means clustering (membership > 0.8), were included in the tree. The colors of the branches correspond to increasing (yellow) or decreasing (blue) trends. The two outer rings represent phyla (first) or genera (second) information. Pie charts reflect the relative proportions of specie within a phylum that increased or decreased over time.
(F) The comparison of the top three abundant phyla in the DI and SI regions. The *p*-values were calculated by the Wilcox test.
(G) PCoA analysis of samples with Bray-Curtis distance, showing that microbiome composition transited along the duration of restoration ($P < 0.001$, adonis). The colors of nodes represent samples of different restoration duration.
(H) Ternary plots of beta diversity comparisons (using the Sørensen dissimilarity index) for different kingdoms. Each point represents a pair of sites and 3204 pairs of sites are included. The position of each dot is determined by a combination of values of the S (similarity), Repl (replacement), and RichDiff (richness difference) matrices. The mean values of S, Repl, and RichDiff are shown below each plot.
(I) Plotting of asymmetric eigenvector maps (AEM) variables over time. AEM1 had the lowest *p*-values and increased over time, suggesting that AEM1 represents the principal temporal trend of microbiota during restoration in DI regions.
(J) Partial dbRDA variation-partitioning analysis.
(K) The Adonis analysis of the forward-selected variables used in the dbRDA analysis.
(L) Results of variation partition with forward-selected variables using dbRDA after grouping variables into categories; these numbers were used to plot the (J).

(M) Ternary plots of variation-partitioning analysis of highly prevalent genera (579 genera in 81 samples) in different domains of life. Each dot represents a genus, and the size of the dot corresponds to the total explained variation. Depending on the genera, either physiochemical (dark yellow) or temporal (blue) variables may play dominant roles, or neither (gray). Contours denote 0.1 to 0.9 confidence intervals.

**Figure S10. The functional diversity of the DI regions and the monoculture site, related to Figures 3 and 4.**

(A) Key characteristics of the species interaction networks in SD control and DI regions. The dynamic characteristics with *p.adj* < 0.05 by GLM in the DI regions are labeled directly.

(B) Community stability in SD control and DI regions. Different lowercase letters above boxplots indicate $p < 0.05$. The GLM statistics are shown for the DI regions.

(C) Heatmap of the differentially abundant KO pathways (*p.adj* < 0.5) based on regression analyses in the DI regions. The functional categories were labeled yellow (increasing) or green (decreasing) on the left.

(D) Fuzzy c-means clustering of the KO pathways based on the relative abundance in the DI regions, with four temporal clusters shown.

(E) Absolute abundance of KO pathways based in the DI regions.

(F) Reporter score bar plot comparing KO pathways enriched in the 4-year group compared with the 9-year group in the DI regions. Pathways with a reporter score greater than 2.5 in at least one group are plotted.

(G) Relative abundance of carbohydrate-active enzymes (CAZYmes) in the DI regions. The statistics by GLM are shown. GHs, glycoside hydrolases; GTs, glycosyltransferases; CEs, carbohydrate esterases; PLs, polysaccharide lyases; AA, auxiliary activities; CBMs, carbohydrate-binding modules.

(H) Effects of the duration of restoration (4 - 9 years) on nutrient cycling pathways (GLM; *p.adj* < 0.05).

(I) Shannon indexes of the microbiota in different domains of life at the monoculture site.

(J) The relative abundance of the top 10 phyla at the monoculture site. Note that BC and NR regions have significant presences of Cyanobacteria.

(K) The top three differentially abundant phyla at the monoculture site.

(L) Heatmap of the differentially enriched KO pathways at the monoculture site. The functional categories of interests were labeled yellow (enriched in the BC group) or green (enriched in the NBC group) on the left. The KO pathways with *p.adj* < 0.05 (Wilcox test) in the NBC and the BC groups were shown.

(M) PCA bi-plot of all KEGG pathways from all samples at the monoculture site. Colored arrows denote the relative importance of contributing features. The colors of points denote the grouping information.

(N) The relative abundance of the top 8 superclass-level metabolites at the monoculture site.

(O) The top ten differentially abundant metabolites between BC and NBC groups. Related parent 1, parent 2, or subclass information were labeled. The Wilcox test was used for BC and NBC comparisons. Only top metabolites enriched in the NBC group with *p*-values <0.02 were shown.

**Figure S11. The gradient boosting regression model, SEM, causal, and evolutionary analysis of desert soil microbiome at the polyculture site, related to Figures 6 and 7.**

(A) The top 20 variables, ranked by relative influences, for predicting the Ks by the gradient boosting regression model. Text color indicates the directions of correlations between variables and Ks as determined by GLM.

(B-C) The impact of restoration duration, irrigation methods, and fungal diversity on carbon (B) and sulfur (C) as estimated by structural equation modeling (SEM). Blue lines indicate negative relationships, and orange lines indicate positive relationships. Paths with non-significant coefficients are presented as gray lines. Fungal diversity is represented by the Shannon index; biomass is represented by DNA concentration. C, carbon; S, sulfur; ***$P < 0.001$; **$P < 0.01$; *$P < 0.05$.

(D) Bar plot showing the top genera involved in more than five mediation linkages.

(E) Sankey diagrams chart showing the mediation analysis results of the *Arthrobacter* (left) and *Geodermatophilus* (right).

(F) Sankey plot illustrating the microbiome's role in decreasing (yellow) particle size through metabolites (left) or directly (right). The bottom right displays metabolites that decrease (yellow) or increase (blue) particle size.

(G) SNP density is correlated with nucleotide diversity (π or pi) density in bacteria and fungi.

(H) Sequencing coverage is correlated with SNP density in bacteria and fungi.

(I) The genes with pN/pS lower than 0.01 present in at least ten samples. Each small circle represents a gene filled based on the frequency of occurrence in samples. One hundred twenty-five genes from 11 species are shown. A larger gray circle-packed gene of the same species. Genes were annotated by both Pfam (gene name; black text) and KEGG pathways (colored circles) and marked by functional annotation (colored dashed circles). The genes without names were represented by "Gene number".

(J) The dynamic pN/pS of biogeochemical cycling genes in the SI regions. The correlation coefficient and P value were calculated by the Spearman method. *G. africanus*: *Geodermatophilus africanus,* G. obscurus: *Geodermatophilus obscurus,* FACHB−61: *Microcoleus* sp. FACHB−61, and IPMA8: *Microcoleus asticus* IPMA8.

**Table S1. The desert soil biophysiochemical metadata and sequencing statistics data.**

**Table S2. The desert soil metabolomics abundance and classification datasets.**
