## Supplementary figures and images for "Progressive community, biogeochemical and evolutionary remodeling of the soil microbiome underpins long-term desert ecosystem restoration"

### Figure S1.jpg

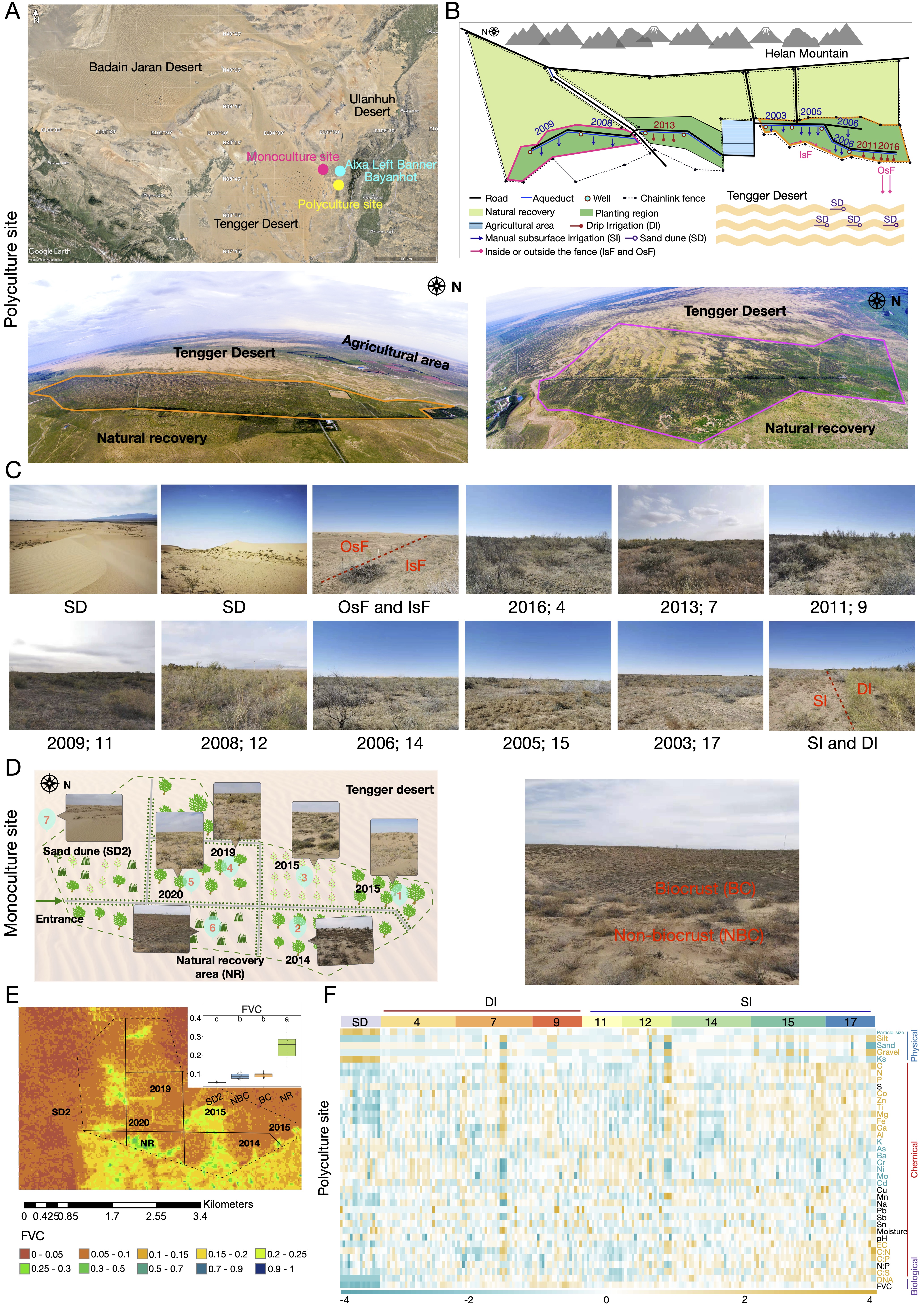

### Figure S2.jpg

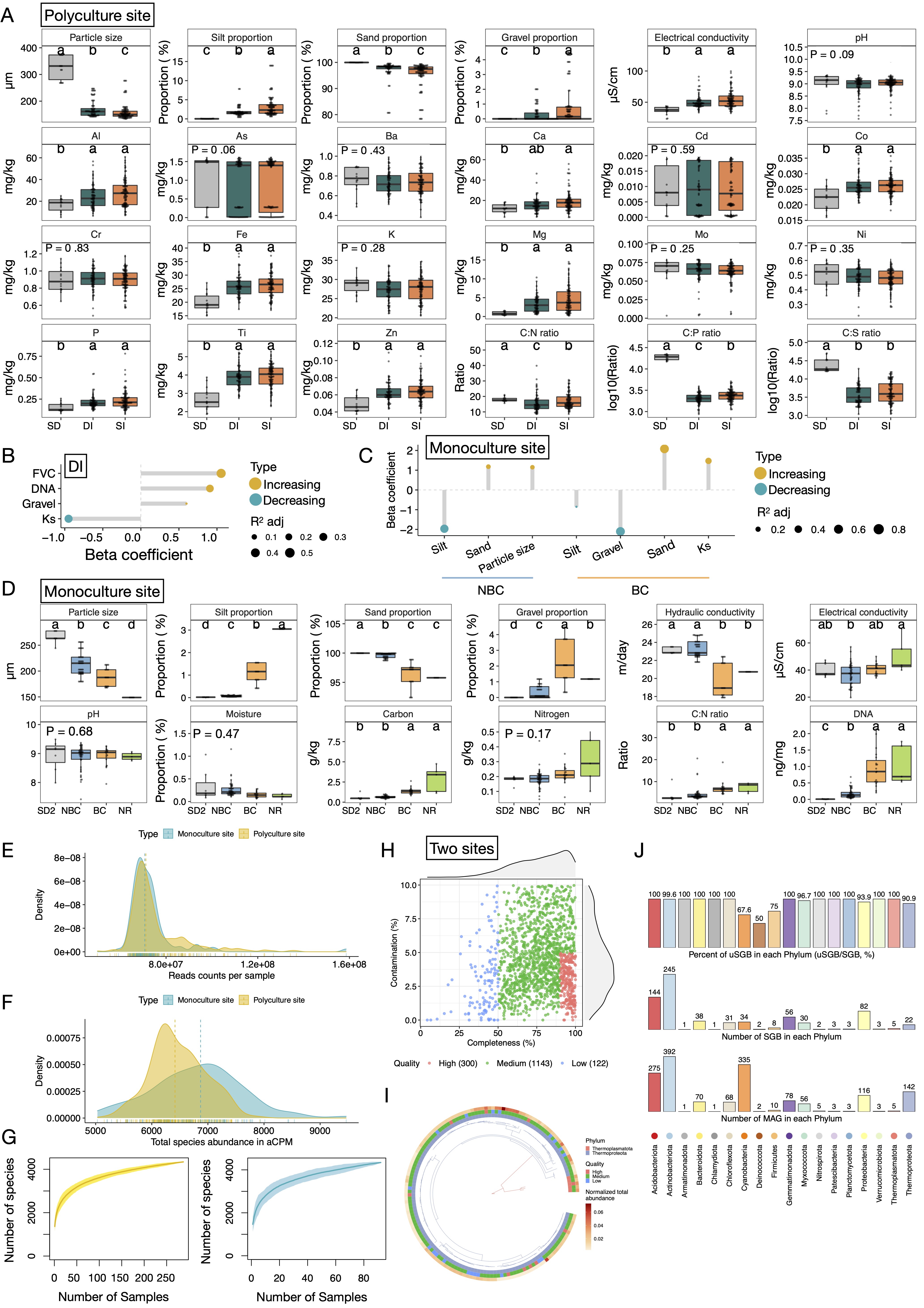

### Figure S3.jpg

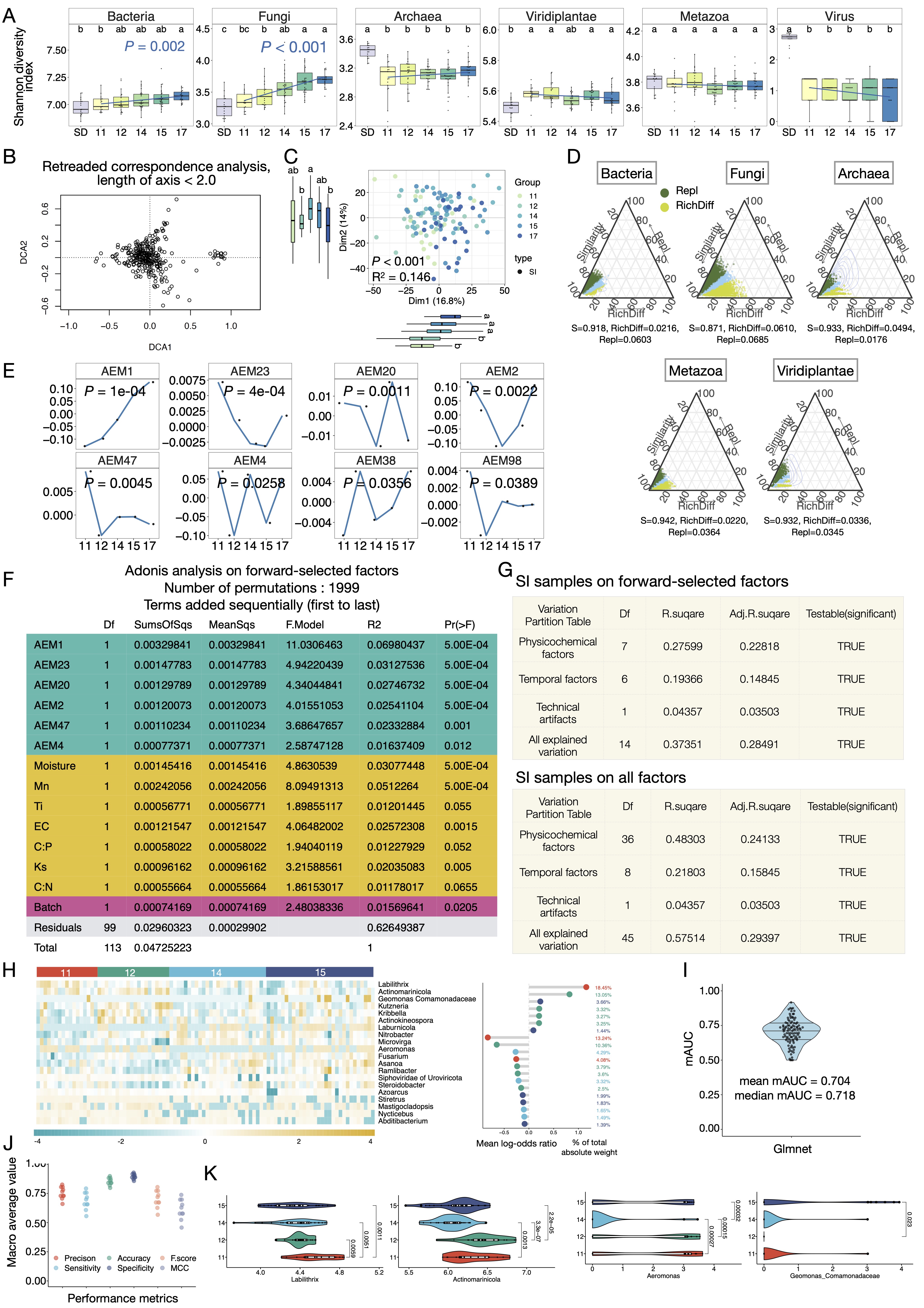

### Figure S4.jpg

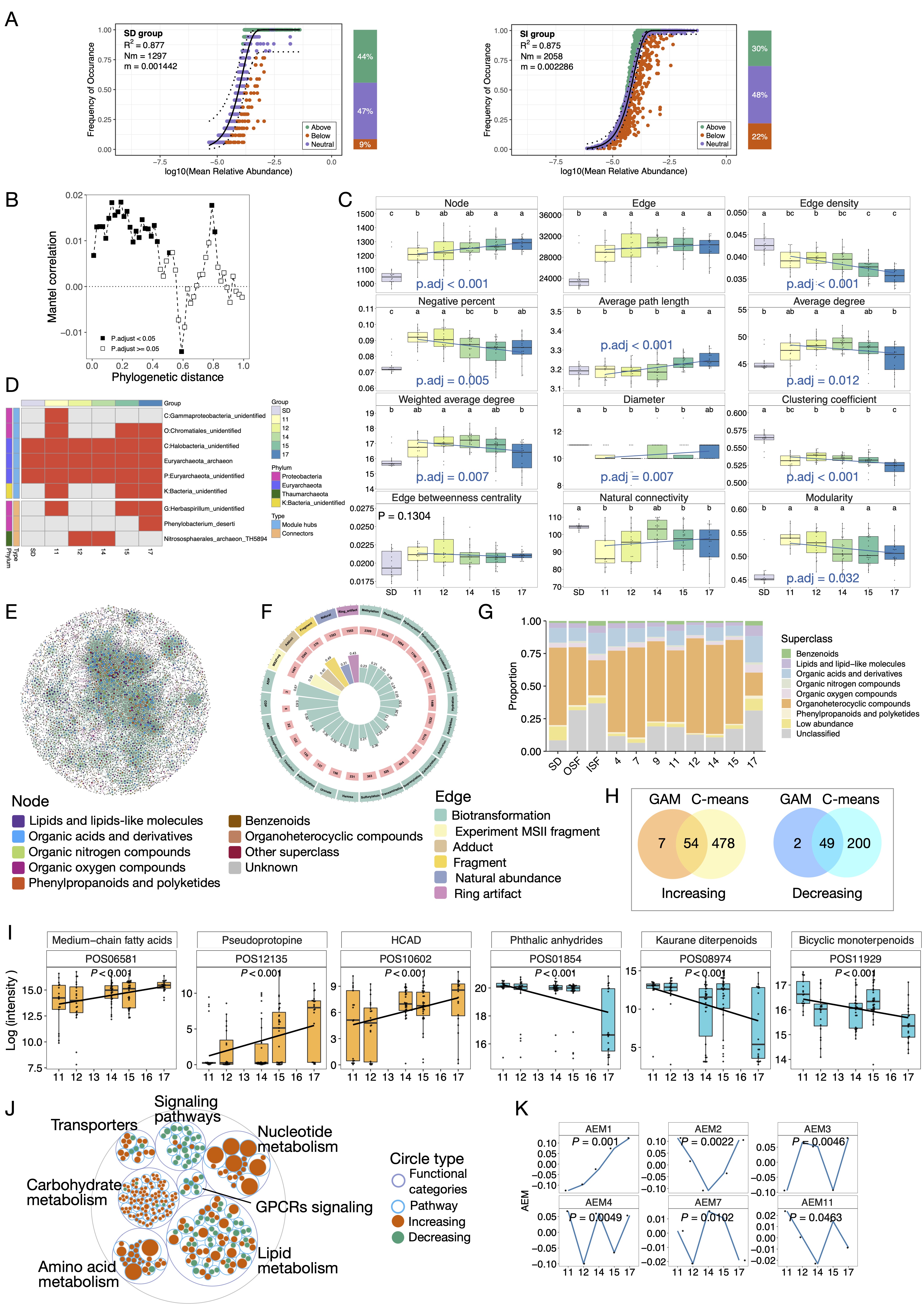

### Figure S5.jpg

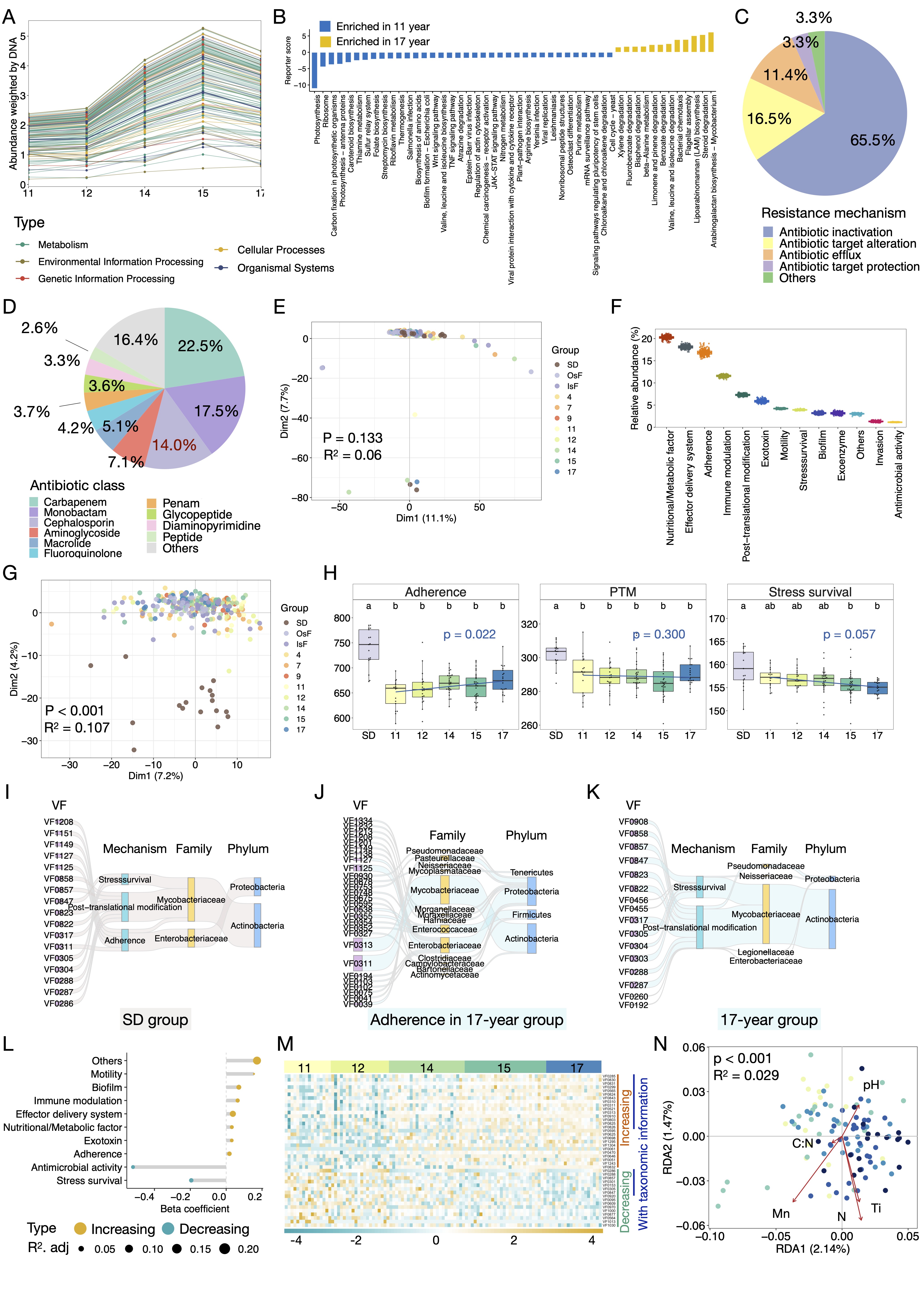

### Figure S6.jpg

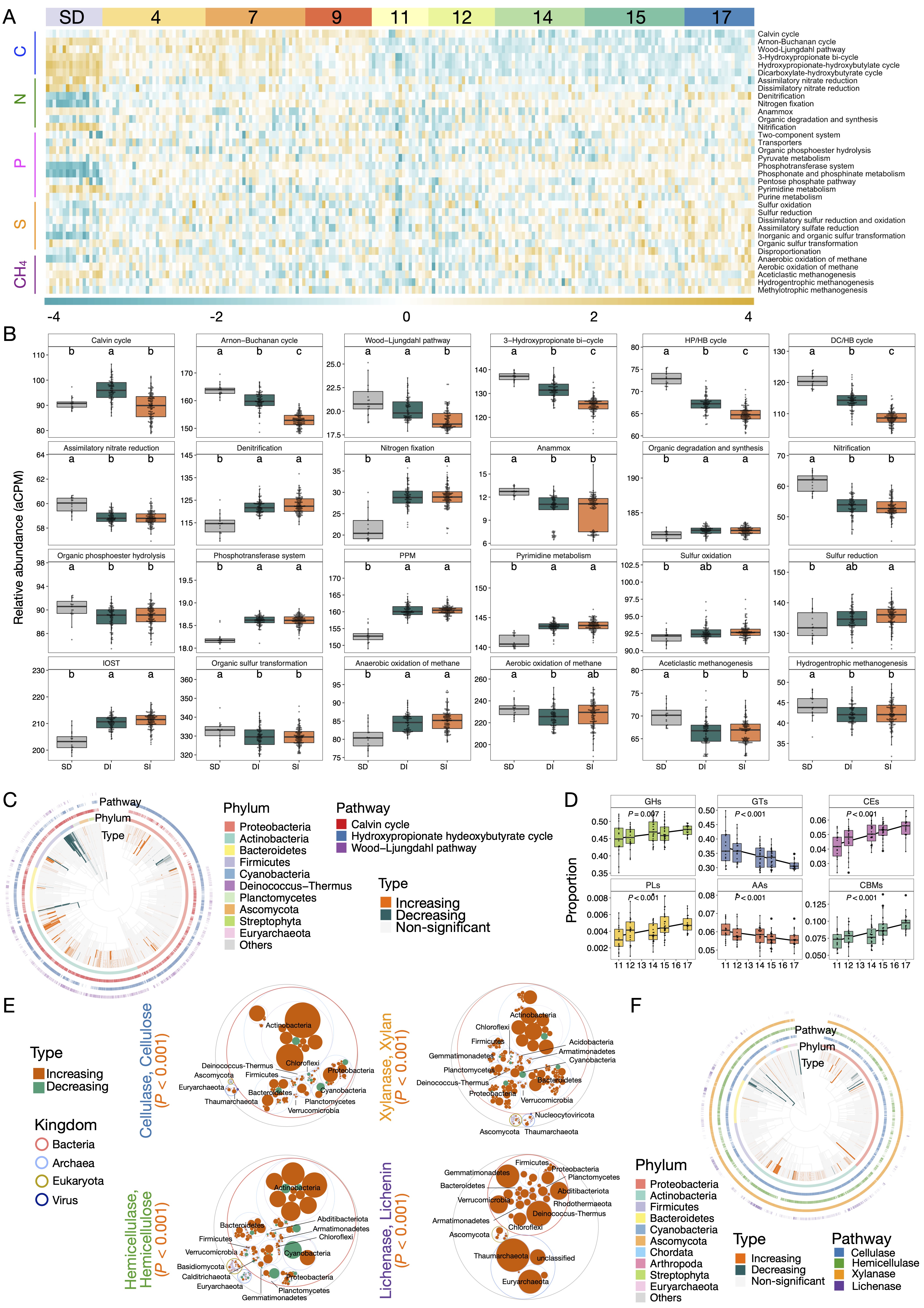

### Figure S7.jpg

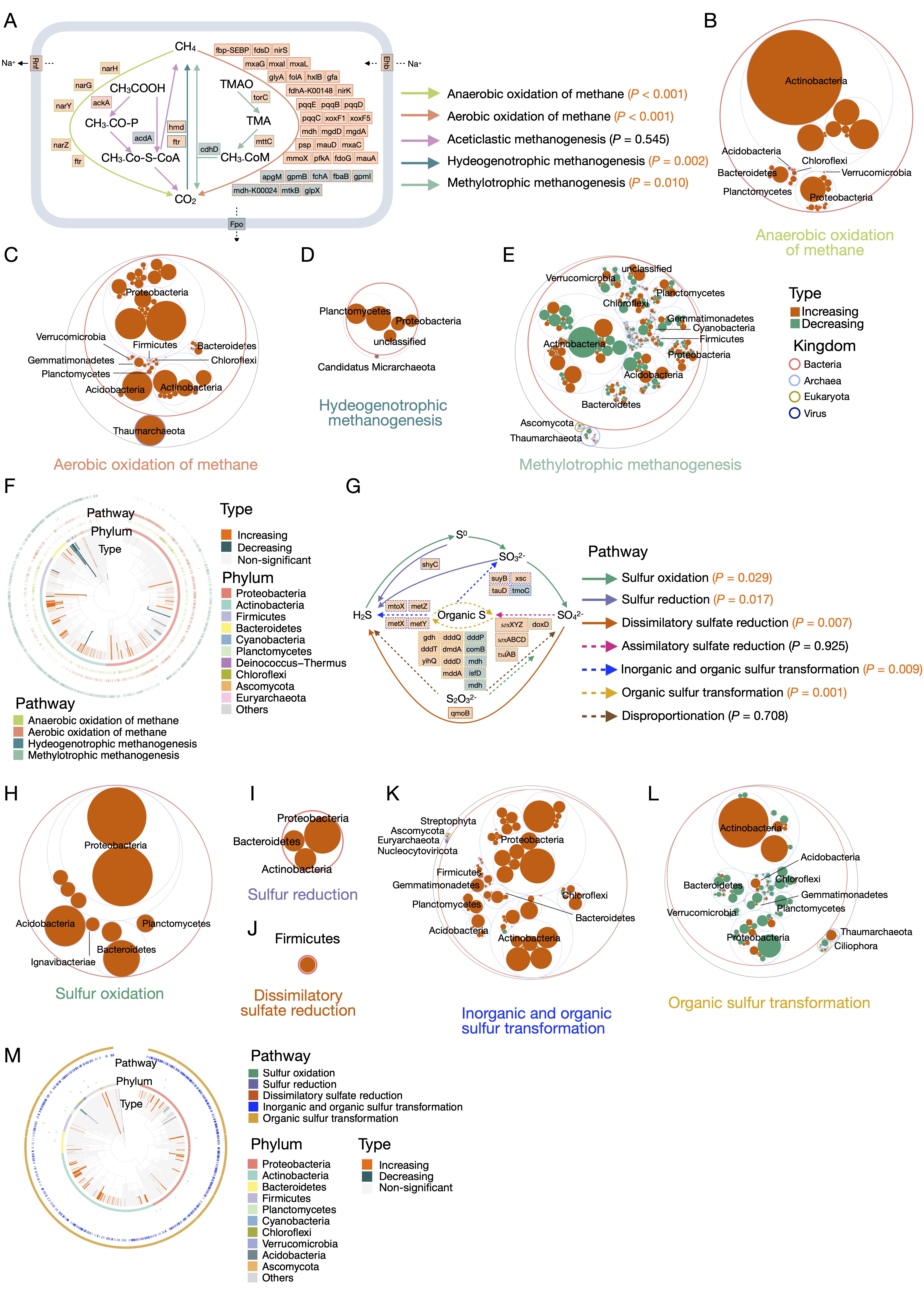

### Figure S8.jpg

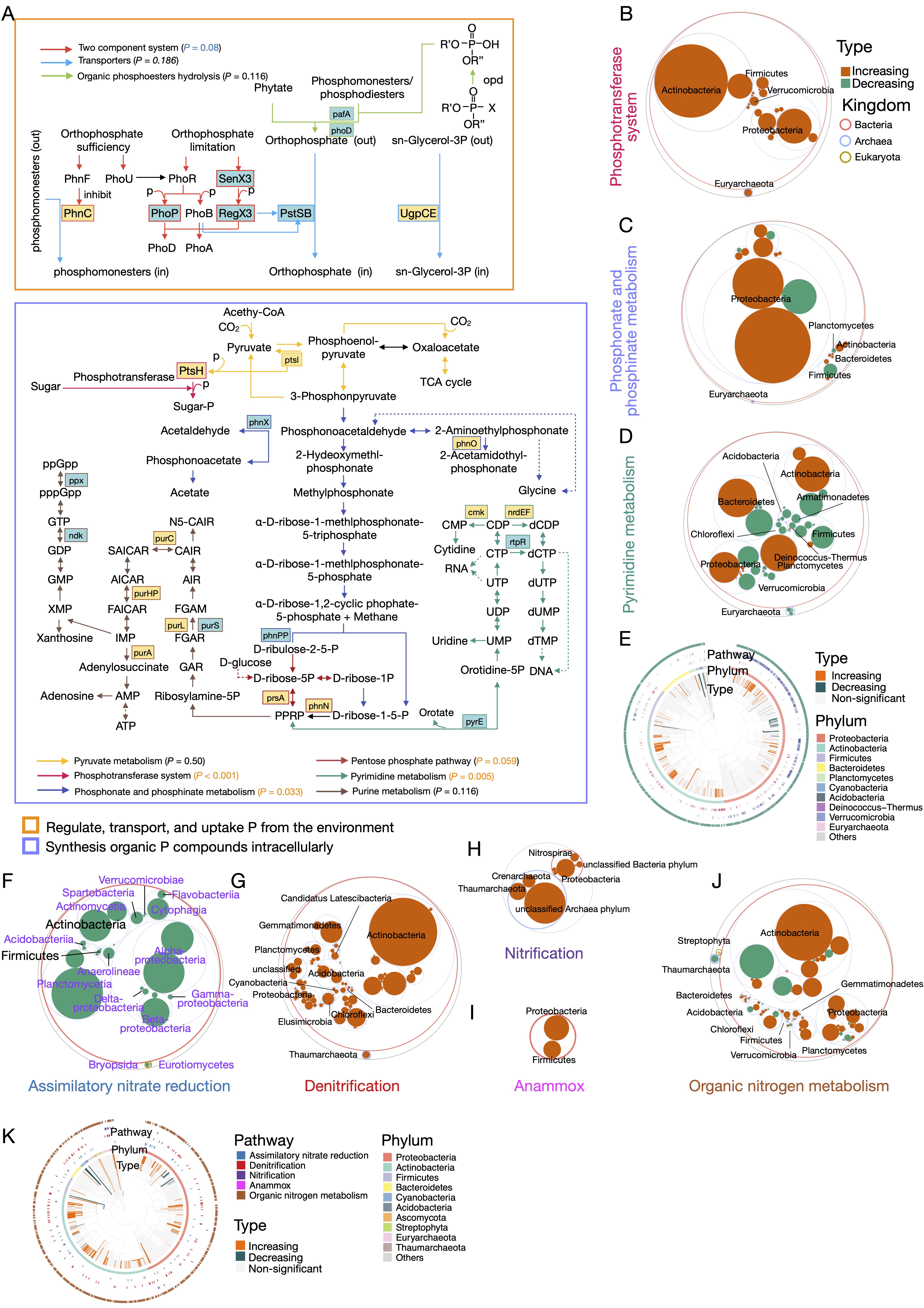

### Figure S9.jpg

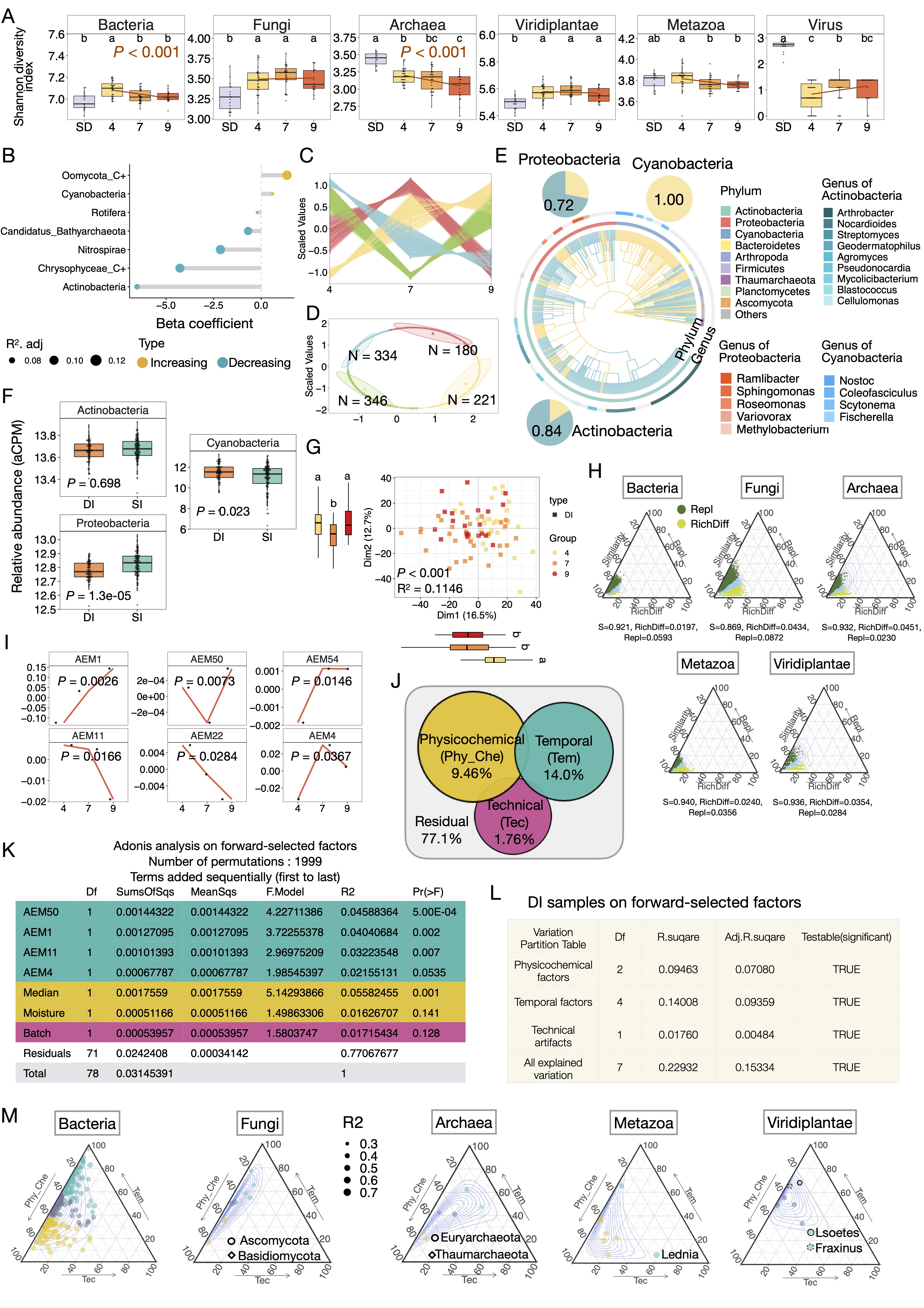

### Figure S10.jpg

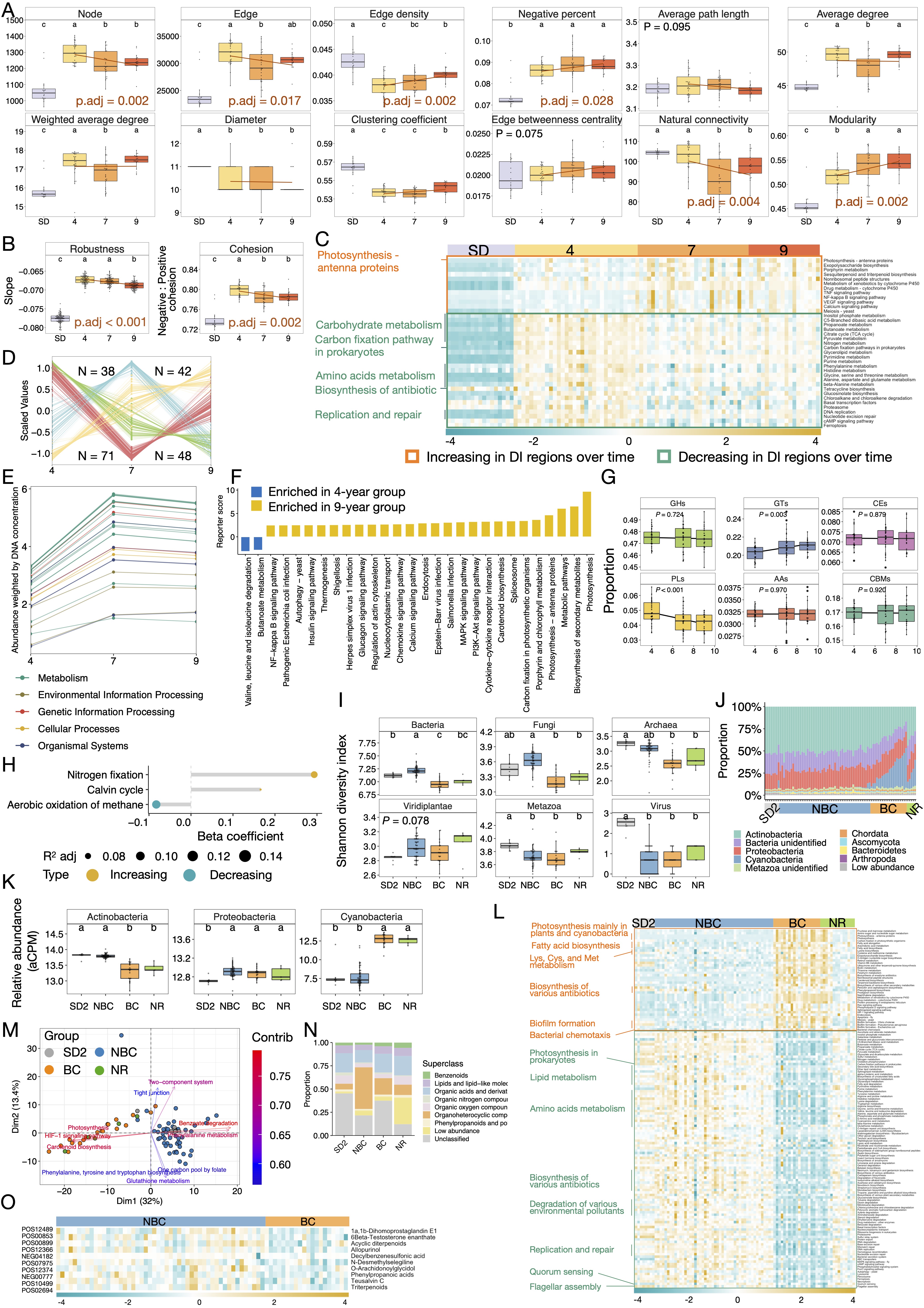

### Figure S11.jpg

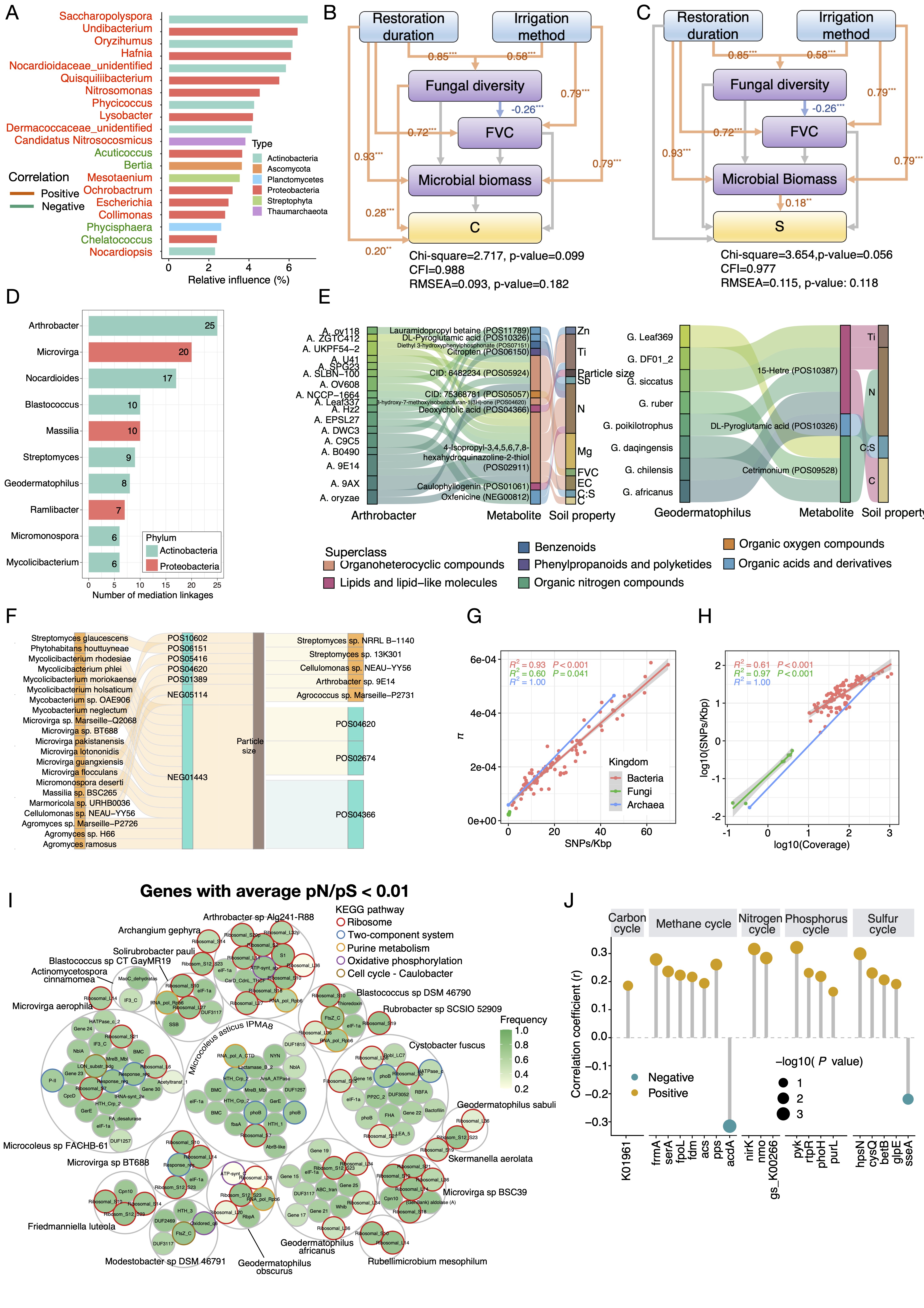
